# A genome-wide analysis of adhesion in *Caulobacter crescentus* identifies new regulatory and biosynthetic components for holdfast assembly

**DOI:** 10.1101/446781

**Authors:** David M. Hershey, Aretha Fiebig, Sean Crosson

## Abstract

Due to their intimate physical interactions with the environment, surface polysaccharides are critical determinants of fitness for bacteria. *Caulobacter crescentus* produces a specialized structure at one of its cell poles called the holdfast that enables attachment to surfaces. Previous studies have shown that the holdfast is a carbohydrate-based material and identified a number of genes required for holdfast development. However, incomplete information about its chemical structure, biosynthetic genes and regulatory principles has limited progress in understanding the mechanism of holdfast synthesis. We have leveraged the adhesive properties of the holdfast to perform a saturating screen for genes affecting attachment to cheesecloth over a multi-day time course. Using similarities in the temporal profiles of mutants in a transposon library, we defined discrete clusters of genes with related effects on cheesecloth colonization. Holdfast synthesis, flagellar motility, type IV pilus assembly and smooth lipopolysaccharide (SLPS) production represented key classes of adhesion determinants. Examining these clusters in detail allowed us to predict and experimentally define the functions of multiple uncharacterized genes in both the holdfast and SLPS pathways. In addition, we showed that the pilus and flagellum control holdfast synthesis separately by modulating the holdfast inhibitor *hfiA.* This study defines a set of genes contributing to adhesion that includes newly discovered genes required for holdfast biosynthesis and attachment. Our data provide evidence that the holdfast contains a complex polysaccharide with at least four monosaccharides in the repeating unit and underscore the central role of cell polarity in mediating attachment of *C. crescentus* to surfaces.

**Importance:** Bacteria routinely encounter biotic and abiotic materials in their surrounding environments, and they often enlist specific behavioral programs to colonize these materials. Adhesion is an early step in colonizing a surface. *Caulobacter crescentus* produces a structure called the holdfast, which allows this organism to attach to and colonize surfaces. To understand how the holdfast is produced, we performed a genome-wide search for genes that contribute to adhesion by selecting for mutants that could not attach to cheesecloth. We discovered complex interactions between genes that mediate surface contact and genes that contribute to holdfast development. Our genetic selection identified what likely represents a comprehensive set of genes required to generate a holdfast, laying the groundwork for a detailed characterization of the enzymes that build this specialized adhesin.

## Introduction

The bacterial cell envelope is a highly dynamic structure that is essential for growth and division (1). Carbohydrate-based compounds often form the outermost layer of the envelope, comprising a specialized surface that each cell displays to the surrounding environment (2). The roles of surface polysaccharides such as capsules, exopolysaccharides and O-antigens in promoting colonization of preferred niches are well established for both free-living and host-associated bacteria (3-5). However, the enzymes that synthesize and export these polysaccharides have been difficult to characterize due to the chemical complexity of the metabolic intermediates (6). Defining the molecular details of how extracellular carbohydrates are produced is critical to understanding bacterial colonization and how it can be controlled.

The aquatic bacterium *Caulobacter crescentus* has a dimorphic lifestyle characterized by an association with exogenous surfaces. Division in *C. crescentus* is asymmetric and produces two distinct cell types, a chemotactic swarmer cell and a sessile, replication-competent stalked cell (7). In response to environmental and developmental signals, the swarmer cell sheds its flagellum, disassembles its pili and transitions into a stalked cell before dividing (8). Stalked cells are named for a specialized envelope extension called the stalk that emerges from the old pole after disassembly of the flagellum and pili. During the swarmer to stalked transition, cells often produce a polysaccharide rich matrix called the holdfast at the site of stalk development (9). This highly adhesive material allows *C. crescentus* to form essentially permanent interactions with exogenous surfaces (Fig 1A) (10).

**Figure 1.**
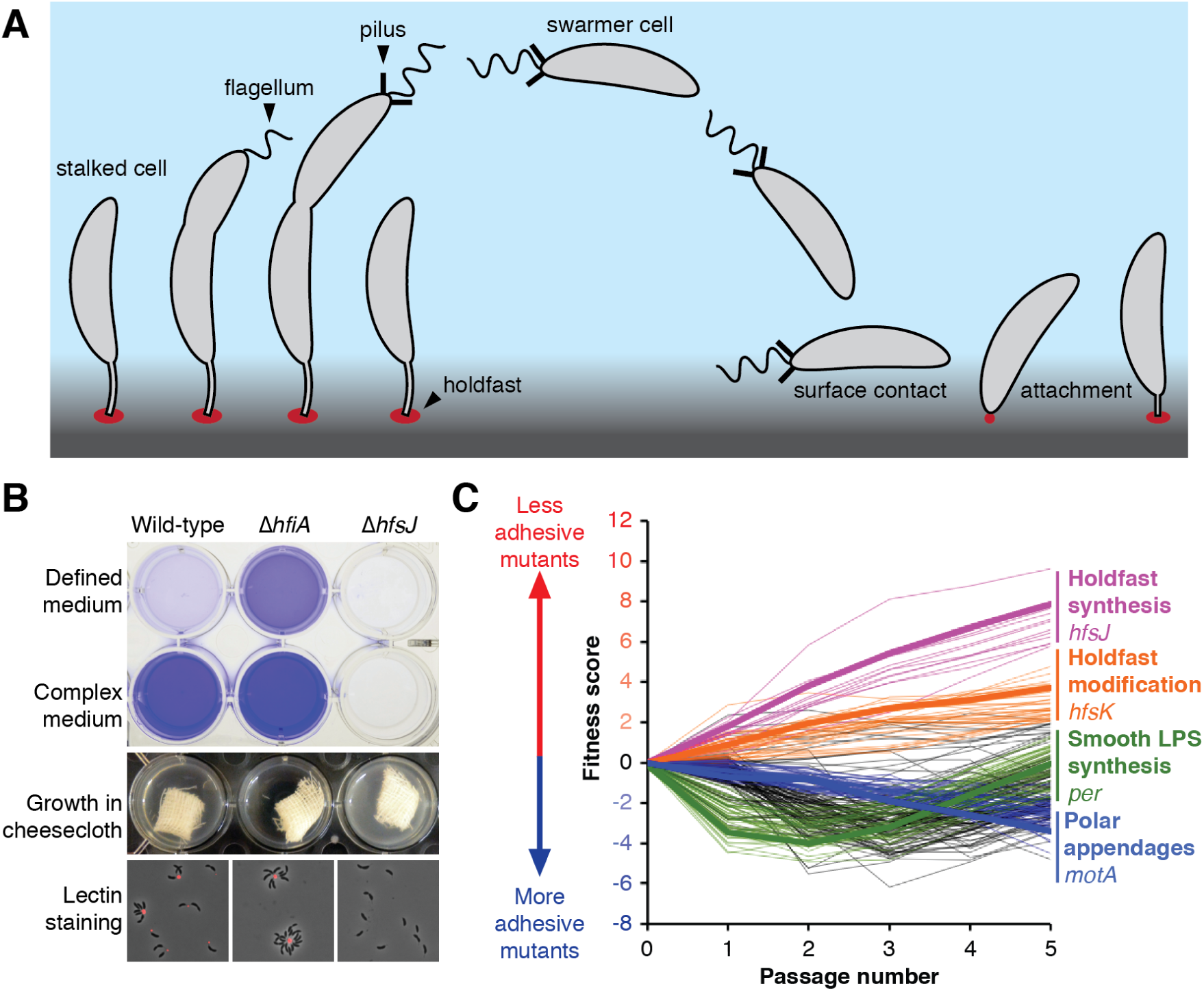
A genome-wide screen for holdfast biosynthesis genes identifies multiple classes of mutants affecting adhesion. A) During the dimorphic *C. crescentus* life-cycle, each cell division produces a motile swarmer cell and a sessile stalked cell. Swarmer cells stop swimming, shed their flagellum and pili and develop into stalked cells before dividing. Stalked cells can adhere strongly to exogenous surfaces using a specialized material called the holdfast. B) The holdfast can be visualized by staining with fluorescently labeled wheat germ agglutinin (fWGA) and biofilm formation can be quantified by crystal violet staining of attached cells. Δ*hfiA* cells overproduce holdfast and are hyper-adhesive. Δ*hfsJ* cells do not produce holdfasts and are non-adhesive. Attachment is reduced in defined medium due to high levels of *hfiA* expression. C) Fitness profiles for the 250 genes with the strongest adhesion phenotypes in the cheesecloth passaging experiment. Lines are drawn to connect mean fitness scores at days zero through five for each of the 250 genes. Genes in the four major fitness clusters are colored according to the legend, with a specific example listed. Genes shown in black do not fit into the any of the four clusters.

Due to the irreversible nature of surface attachment in *C. crescentus*, the timing of holdfast production is tightly controlled. When grown in defined medium only a small proportion of cells produces a holdfast, as compared to nearly all cells when grown in complex medium (11). This effect is due to elevated expression of the *<u>h</u>oldfast inhibitor <u>A</u>* (*hfiA*) gene in defined medium (Fig 1B). *hfiA* expression is also coordinated with the cell-cycle. Its transcript levels drop during the swarmer to stalked transition, which corresponds to the developmental stage at which holdfasts begin to appear. Numerous signaling pathways target the *hfiA* promoter, allowing the cell to integrate environmental, nutritional and developmental cues into a single output that regulates adhesion (11, 12). In yet another regulatory regime, holdfast synthesis can be induced when a swarmer cell encounters a surface (13). Physical disruption of flagellar rotation or pilus retraction upon surface contact stimulates the production of a holdfast (14, 15). How the numerous regulatory pathways converge to control holdfast development remains unclear, but the complexity of these networks reflects the significance of committing to a surface-associated lifestyle.

Genetic analysis of non-adhesive mutants indicates that the holdfast is a polysaccharide-based material. The <u>h</u>oldfast <u>s</u>ynthesis (*hfs*) genes include predicted glycosyltransferases, carbohydrate modification factors and components of a *wzy-*type polysaccharide assembly pathway (16-19). *wzy*-dependent carbohydrate assembly utilizes a lipid carrier known as undecaprenylpyrophosphate (UPP) on which glycosyltransferases assemble an oligosaccharide repeating unit in the cytoplasm (20). The resulting glycolipid is flipped from the cytoplasmic face of the inner membrane to the periplasmic face where the oligosaccharide is polymerized and exported to the cell surface (21). The *wzy* mechanism is used to produce an impressive diversity of polysaccharides and is broadly conserved among bacteria (22). Thus, characterizing enzymes involved in the biosynthesis of the holdfast has the potential to uncover broadly applicable principles about how bacteria produce carbohydrate polymers.

The chemical nature of the holdfast matrix remains poorly characterized. The holdfast binds to the N-acetylglucosamine (GlcNAc)-specific lectin wheat germ agglutinin (WGA) and is sensitive to the GlcNAc specific hydrolases chitinase and lysozyme, indicating that GlcNAc is a component of the matrix (23). Little other information about the carbohydrate content has been reported. Extracellular DNA and unidentified protein component(s) contribute to the stiffness of the holdfast, but only mutations in polysaccharide biosynthesis genes or pleiotropic regulators of cell polarity abolish holdfast production (24). Currently, three glycosyltransferase steps are known to be required for holdfast synthesis. An initial reaction carried out by one of the genetically redundant HfsE, PssY or PssZ enzymes is thought to be followed by the actions of HfsG and HfsJ, suggesting that the polysaccharide may be composed of a triscaccharide repeat (11, 18). However, new *hfs* genes continue to be discovered, hinting that additional glycosyltranferases may remain unidentified (11, 25). Uncertainty about both the composition of the holdfast and the saturation of screens for *hfs* genes presents a major obstacle to characterizing enzymatic reactions in the pathway.

Here, we utilized saturating transposon mutagenesis to probe holdfast production at the genome scale. We developed a barcoded transposon library in *C. crescentus* and enriched for non-adhesive mutants by passaging across multiple days in presence of cheesecloth. We discovered a surprising number of genes with distinct adhesion phenotypes that ranged from hyper-adhesive to non-adhesive. We found that disrupting the smooth lipopolysaccharide (SLPS) leads to a holdfast-independent form of ectopic adhesion that is not restricted to the cell pole but rather mediated throughout the cell surface. The temporal adhesion profiles of known SLPS mutants were used to identify and characterize new genes in the SLPS pathway. The same fitness correlation approach was used to place previously uncharacterized genes in the holdfast pathway. We further demonstrated that disrupting the assembly of polar surface appendages modulates the activity of the holdfast inhibitor, *hfiA.* In particular, individual mutations in the pilus machinery had a range of adhesion phenotypes suggesting that distinct intermediates in the pilus assembly pathway have opposing effects on *hfiA*. Based on our comprehensive analysis of holdfast regulation, biosynthesis and assembly, we propose a model that outlines the sequence of enzymatic steps required to produce the holdfast polysaccharide.

## Results

### A screen for mutants with altered adhesion

The holdfast promotes adhesion of *C. crescentus* cells to a variety of surfaces (26). We reasoned that adhesive cells could be depleted from liquid cultures by adding an attachment substrate with a sufficiently large surface area. Cheesecloth has been used in this manner to enrich for holdfast mutants in both *C. crescentus* and *Asticcacaulis biprosthecum,* another stalked bacterium in the *Caulobacteraceae* clade (27, 28). Adding sterile cheesecloth to wild-type *C. crescentus* cultures decreased the turbidity of the medium by titrating adhesive cells from the broth. This effect was amplified in the hyper-adhesive Δ*hfiA* strain and not observed in the holdfast deficient Δ*hfsJ* strain demonstrating the effectiveness of cheesecloth at capturing cells with a holdfast (Fig 1B). We concluded that growth in the presence of cheesecloth could be used as the basis of a selection to identify mutants defective in adhesion.

Saturating transposon mutagenesis coupled with transposon insertion sequencing (TnSeq) offers the advantage of scoring phenotypes for all non-essential genes in the genome simultaneously (29). Thus, combining TnSeq-based mutant profiling with cheesecloth depletion seemed appropriate to perform a saturating screen for holdfast biosynthesis genes and identify missing biosynthesis factors. We developed a randomly barcoded transposon library in *C. crescentus* to enable the use of BarSeq (30) for profiling mutant fitness. Adhesive cells were depleted by passaging the library in cheesecloth for five cycles. During each passage, the library was cultured for 24 h in the presence of cheesecloth after which the unattached cells in the medium were used to re-inoculate a fresh culture containing cheesecloth. An aliquot of unattached cells in the medium was also harvested for BarSeq analysis. Three passaging experiments with cheesecloth were performed in parallel. To discriminate mutants with adhesion defects from those with growth defects, we also performed three passaging experiments without cheesecloth for comparison (Table S1).

The abundance of each mutant during the passaging steps was assessed using BarSeq, providing a temporal fitness profile for each gene over the course of the experiment (30). Genes with positive fitness scores reflect mutants with adhesion defects that were enriched in medium that had been depleted with cheesecloth. Genes with negative fitness scores represent hyper-adhesive mutants that are depleted more efficiently by cheesecloth than wild type. As expected most genes had inconsequential effects on adhesion. However there were a significant number of genes whose mutation caused strong cheesecloth dependent changes in abundance over the course of the multi-day experiment. The 250 genes with the highest fitness values, above or below the baseline level, are shown in Fig 1C. The time resolved nature of the experiment allowed us to group mutants with similar fitness profiles into distinct classes. Mutants with similar temporal adhesion profiles often mapped to genes with similar or complementary annotations, indicating that they represented groups of functionally related genes. We predicted the functions of uncharacterized genes using known functions for genes with similar fitness profiles. For example, a cluster of genes whose mutants show a continuous increase in abundance after each passage contains many known *hfs* genes, and any uncharacterized genes that share this fitness profile would be predicted to contribute to holdfast synthesis as well. We identified four clusters containing mutants that display distinct fitness profiles for the cheesecloth passaging experiment. Each cluster is described in detail below.

### Mutants defective in smooth lipopolysaccharide display ectopic adhesion

We identified a cluster of mutants with strong fitness decreases (i.e. increased adhesion to cheesecloth) in early passages with relative abundances that recovered to near neutral or even positive fitness values as passaging proceeded (Fig 2A). Many of the genes in this “recovery” cluster had annotations associated with polysaccharide biosynthesis but had no known cellular function. However, the *wbq* genes that are required for the production of smooth lipopolysaccharide (SLPS) comprised a subset of the recovery cluster (31). This suggested that mutants sharing this fitness profile might also be defective in the biosynthesis of SLPS.

**Figure 2.**
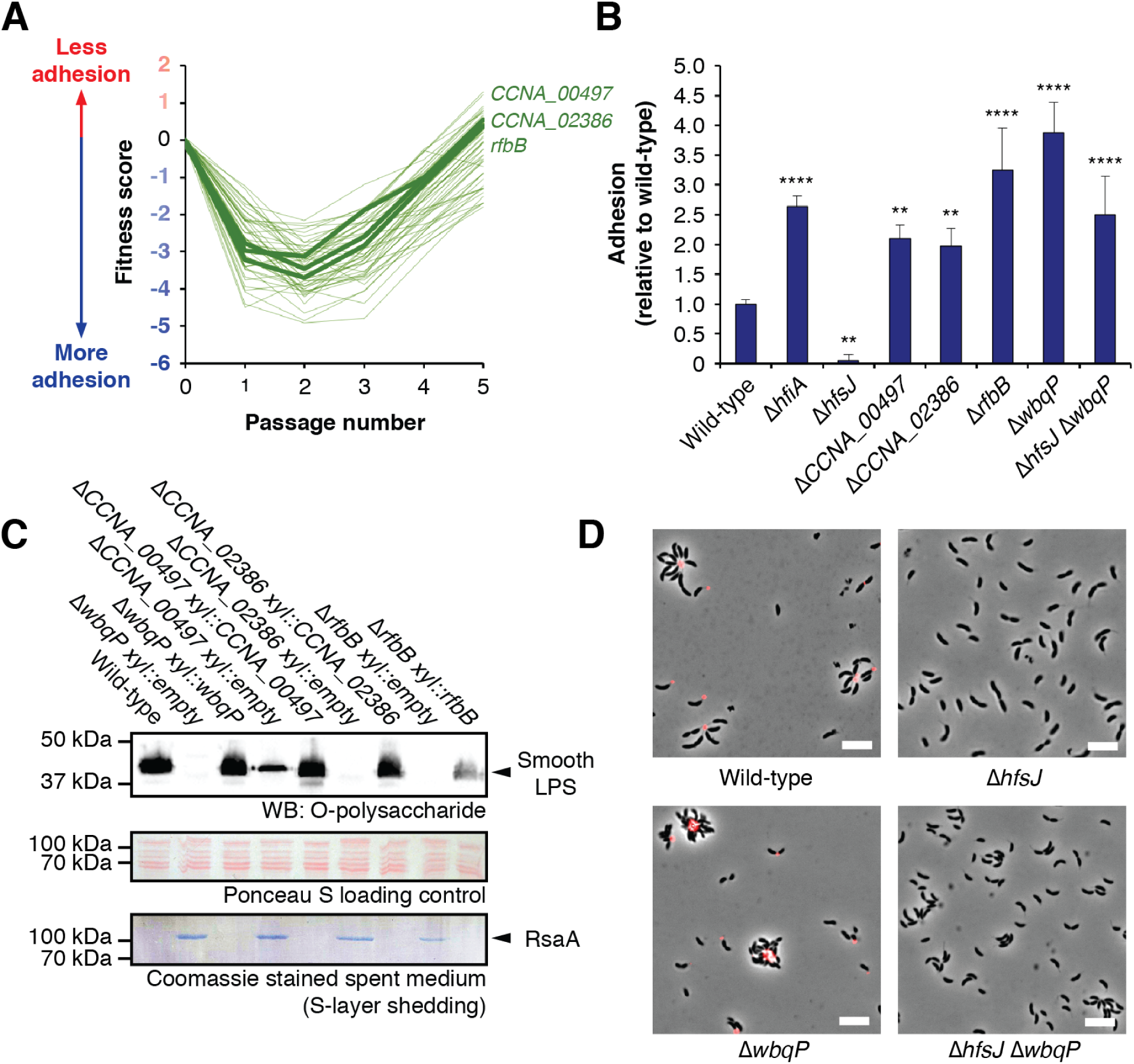
Disrupting SLPS production leads to ectopic adhesion. A) Fitness profiles for genes in the SLPS cluster. A full list of these genes and their annotations is provided in Table S5. *wbqP* was not characterized in our library due to low insertion density in this region. B) Surface attachment of SLPS mutants measured by CV staining. Cultures were grown for 24 hours in M2X medium before staining surface attached cells. Disrupting SLPS leads to increased adhesion in a holdfast-independent fashion. The graph shows the average (± standard deviation) of five biological replicates. Statistical significance was assessed by ANOVA with a pairwise Dunnett’s post test to determine which samples differed from wild type. ** *P* < 0.01; **** *P* < 0.0001. C) Smooth LPS production is disrupted in Δ*CCNA_00497,* Δ*CCNA_02386* and Δ*rfbB*. Top: western blot to detect SLPS. Middle: Total protein stained with ponceau S as a loading control for SLPS blot. Bottom: Coomassie staining of spent medium from the cultures. Each mutant shows a loss of or a decrease in SLPS production by western blot and releases the S-layer protein RsaA into the spent medium. The cell-surface defects can be complemented by ectopic expression of the appropriate gene. “Empty” refers to plasmid control strains

We focused on three uncharacterized genes in the recovery cluster. *CCNA_00497* is annotated as a putative rhamnosyl transferase, *CCNA_02386* is annotated as an O-antigen ligase and *CCNA_03744* is homologous to *rfbB*, a gene required for the biosynthesis of dTDP-L-rhamnose (32). Mutations in *rfbB* were previously shown to suppress the holdfast attachment defect observed in a Δ*hfaD* mutant, but SLPS was not examined in these mutants (33). We created in-frame deletions of *CCNA_00497, CCNA_02386* and *rfbB* and analyzed SLPS production by immunoblotting. A deletion of *wbqP*, which is thought to encode the initial glycosyltransferase step in the O-polysaccharide biosynthesis pathway, was used as a positive control. Disruption of *CCNA_02386, rfbB* or *wbqP* led to the loss of detecTable SLPS, and Δ*CCNA_00497* cells showed a reduction in SLPS levels (Fig 2C). Additionally, all four mutants released the S-layer protein RsaA into the spent medium, an additional hallmark of SLPS defects in *C. crescentus* (Fig 2C) (34). None of the mutants displayed observable changes in rough LPS, demonstrating that they were not defective in the production of lipid A or the core oligosaccharide (Fig S1). All of the defects could be complemented by ectopic expression of the target gene, confirming their roles in the production of SLPS (Fig 2C and Table S2).

The fitness profiles for early stages of cheesecloth passaging suggested that disrupting SLPS led to hyper-adhesive cells that were rapidly depleted by cheesecloth. In complex medium, nearly all wild-type cells produce a holdfast, making the dynamic range for detecting increased adhesion quite small. Thus, we chose to investigate potential hyper-adhesive phenotypes by examining mutants using a defined medium in which fewer cells produce a holdfast. We found that the SLPS mutants were indeed hyper-adhesive, producing CV staining values ranging from two to four times that of wild type (Fig 2B). To understand the basis of hyper-adhesion, Δ*wbqP* cells were imaged after staining with fluorescently labeled wheat germ agglutinin (fWGA) to label holdfasts. Most wild-type cells displayed a fluorescent focus at the tip of the stalk, and stalks from multiple cells often aggregated around a single focus to form rosette structures that are characteristic of holdfast production. In the Δ*wbqP* background, a comparable number of cells produced a holdfast, but the structure of the rosettes was altered. Cells that assembled around a holdfast were more tightly packed, and not all of them adhered to the aggregates through the tip of the stalk (Fig 2D).

The unusual rosette structures in the Δ*wbqP* mutant suggested that cells with disrupted SLPS might have a second mode of adhesion that did not require holdfast. We compared fWGA staining in the holdfast deficient Δ*hfsJ* strain to a Δ*hfsJ* Δ*wbqP* double mutant that lacks both holdfast and SLPS. Δ*hfsJ* cells did not stain with fWGA and did not form aggregates. Δ*hfsJ* Δ*wbqP* cells did not stain with fWGA, but in contrast to the Δ*hfsJ* strain, the cells still formed aggregates (Fig 2D). These aggregates appeared not to be mediated by stalk-stalk interactions but rather through interactions of the cell body. This further supported the idea that a holdfast-independent mode of ectopic adhesion operates in SLPS mutants. Consistent with this model, bulk adhesion in the Δ*hfsJ* Δ*wbqP* double mutant was not abolished, and was, in fact, higher than wild type (Fig 2B). We conclude that disrupting SLPS production causes defects in the cell surface leading to a holdfast independent mode of adhesion that represents the dominant mode of adhesion for these mutants early in our experimental time course.

### The flagellum and type IV pili regulate holdfast production

A second cluster of mutants primarily contained genes known to participate in chemotaxis and flagellar motility as well as components of the type IV pilus machinery. The fitness profiles for these mutants suggested that disrupting the assembly of polar appendages, either pili or the flagellum, leads to hyper-adhesion. To study the effects of polar appendages on adhesion, we deleted the genes for the flagellar basal body component FlgH and the pilus assembly protein CpaH (35, 36). We confirmed that Δ*flgH* showed the expected loss in motility and that Δ*cpaH* was resistant to the type IV pilus specific phage ΦCBK (Figs S2 and S3).

Both the Δ*flgH* and the Δ*cpaH* mutants showed adhesion defects in complex medium (Table S2). However, in defined medium, both mutants displayed increased adhesion relative to wild type, indicating that disrupting the pilus or the flagellum causes hyper-adhesion under these conditions (Fig 3B). To reconcile these differences we used fWGA staining to measure the proportion of cells that produced a holdfast. The Δ*flgH* and Δ*cpaH* mutants produced more holdfasts than wild-type in both complex and defined medium (Fig S3 and Table S3). We conclude that flagellum and pilus mutations increase holdfast production, but that loss of either appendage also leads to holdfast-independent defects in surface colonization. Pili and flagella often have similar effects on surface colonization in other systems (37, 38). Because the baseline level of holdfast production is low in defined medium, the enhanced holdfast production Δ*flgH* and Δ*cpaH* backgrounds appears to outweigh surface colonization defects under these conditions (Fig S3).

**Figure 3.**
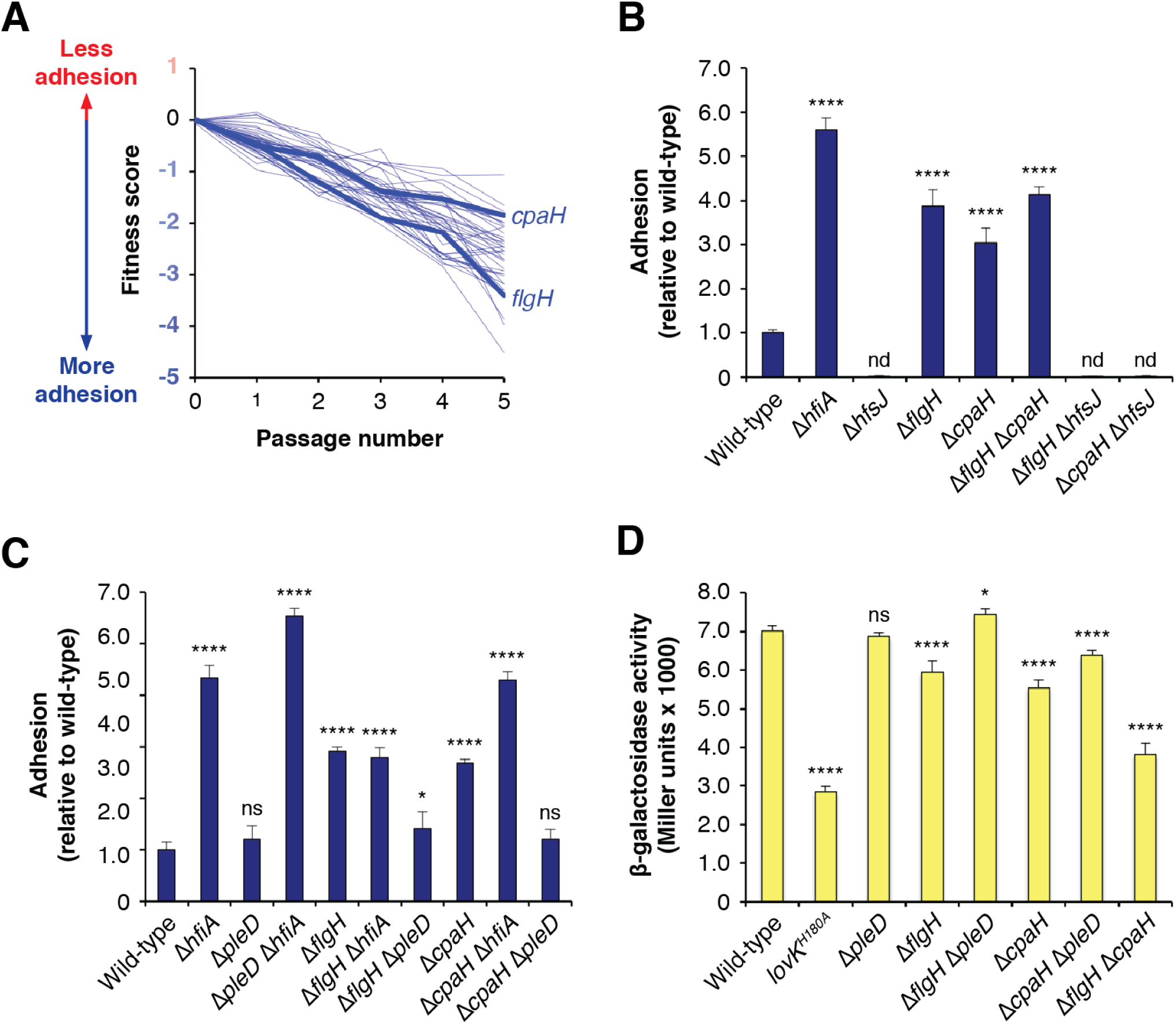
Disrupting polar appendages stimulates holdfast production. A) Fitness profiles for genes in the polar appendage cluster. A full list of these genes and their annotations is provided in Table S5. B) Surface attachment of motility mutants measured by CV staining. Cultures were grown for 17 hours in M2X medium before staining surface attached cells. Deletion of the genes for either the outer-membrane flagellar base protein FlgH or the inner-membrane type IV pilus component CpaH caused increased adhesion. In both mutant backgrounds, *hfsJ* is required for attachment. The graph shows the average (± standard deviation) of five biological replicates. C) Effect of *hfiA* and *pleD* deletions on surface attachment of *cpaH* and *flgH* mutants. Δ*hfiA* does not affect adhesion in the Δ*flgH* background and increases adhesion in the Δ*cpaH* background. The increased adhesion in both the Δ*flgH* and Δ*cpaH* mutants can be eliminated by deletion of *pleD.* The graph shows the average (± standard deviation) of four biological replicates. D) *P*_*hfiA*_*-lacZ* reporter activity in polar appendage mutants. The chart shows the average (± standard deviation) of four biological replicates. Statistical significance was assessed by ANOVA with a pairwise Dunnett’s post test to determine which samples differed from wild type. nd – not detected; ns – not significant; * *P* < 0.05; **** *P* < 0.0001. When necessary, P values for additional pairwise comparisons pertinent to interpretation are indicated in the text.

A recent report showed that flagellar hook mutants displayed decreased transcription from the *hfiA* promoter (*P*_*hfiA*_) in defined medium and that this effect did not occur in the absence of the pleiotropic cell-cycle regulator PleD (39-41). This led us to examine the relationships between our polar appendage mutants, *hfiA* and *pleD*. We used a *P*_*hfiA*_*-lacZ* reporter to measure transcription from the *hfiA* promoter. Because expression from *P*_*hfiA*_ is low in complex medium, the dynamic range for measuring decreased activity is small. Therefore, we focused on transcriptional changes that occurred in defined medium where the baseline activity of *P*_*hfiA*_ is high. The Δ*flgH* and Δ*cpaH* mutants showed reduced *hfiA* transcription in the reporter assay (Fig 3D). This modest reduction in *P*_*hfiA*_*-lacZ* reporter signal is statistically significant, and changes of this magnitude are known to affect holdfast development (11). The decrease in *P*_*hfiA*_*-lacZ* signal was abrogated in the Δ*flgH* Δ*pleD* and Δ*cpaH* Δ*pleD* double mutants (Fig 3D). Likewise, bulk adhesion in defined medium reverted to near wild-type levels in the Δ*flgH* Δ*pleD* and Δ*cpaH* Δ*pleD* mutants, confirming that *pleD* contributes to the modulation of *P*_*hfiA*_ in the pilus and flagellar mutants (Fig 3C). We note, however, that a full reversion of the hyper-holdfast phenotype would be predicted to display bulk adhesion levels below that of wild type due to the holdfast-independent attachment defects seen in pilus and flagellar mutant backgrounds. Thus, while *pleD* does contribute to the enhanced adhesion seen in the Δ*flgH* and Δ*cpaH* mutants, the effect is not completely dependent on this gene.

To test whether the hyper-adhesive phenotypes in the polar appendage mutants could be explained by repression of *hfiA,* we created Δ*flgH* Δ*hfiA* and Δ*cpaH* Δ*hfiA* double mutants. Bulk attachment in the Δ*flgH* Δ*hfiA* strain was not significantly increased relative to the Δ*flgH* single deletion (Fig 3C). This suggests that Δ*flgH* effectively inactivates the effects of *hfiA*, and that holdfast-independent adhesion defects lower the maximum level of surface attachment that can be achieved in a Δ*flgH* background. The enhanced attachment seen in the Δ*cpaH* background was further increased in a Δ*cpaH* Δ*hfiA* double mutant (*P* < 0.0001; Fig 3C). Thus, Δ*cpaH* has an intermediate effect on *hfiA* activity by dampening but not completely masking its activity. Consistent with this, the fraction of Δ*cpaH* cells that produced a holdfast in defined medium was intermediate between that of wild type and Δ*hfiA*, supporting the idea that Δ*cpaH* causes both intermediate enhancement of holdfast production and holdfast-independent surface-attachment defects (Fig S3 and Table S3). Finally, bulk adhesion in a Δ*flgH* Δ*cpaH* double mutant was indistinguishable from Δ*flgH*, and *P*_*hfiA*_ transcription was lower in Δ*flgH* Δ*cpaH* than either the Δ*flgH* (*P* < 0.0001) or Δ*cpaH* (*P* < 0.0001) single mutants (Fig 3B and C). These results indicate that holdfast production is likely maximized in the Δ*flgH* mutant and that the Δ*flgH* and Δ*cpaH* mutants modulate *P*_*hfiA*_ through separate pathways.

### A complex role for the pilus in regulating adhesion

The Δ*cpaH* phenotype suggested that disrupting pilus assembly leads to increased holdfast production via the repression of *hfiA*. However, a closer examination of the fitness profiles for genes involved in type IV pilus assembly revealed a range of phenotypes for various components of the apparatus (Fig 4A). Most of the genes encoding components of the pilus secretion machinery, including *cpaH,* had fitness profiles consistent with increased adhesion. However, mutations in the gene coding for the main pilin subunit, PilA, displayed the opposite trend. *pilA* mutants had fitness profiles that would be expected for mutants with adhesion defects. We confirmed that, indeed, the Δ*pilA* strain was defective in surface attachment both in complex and defined medium (Fig 4B and Table S2).

**Figure 4.**
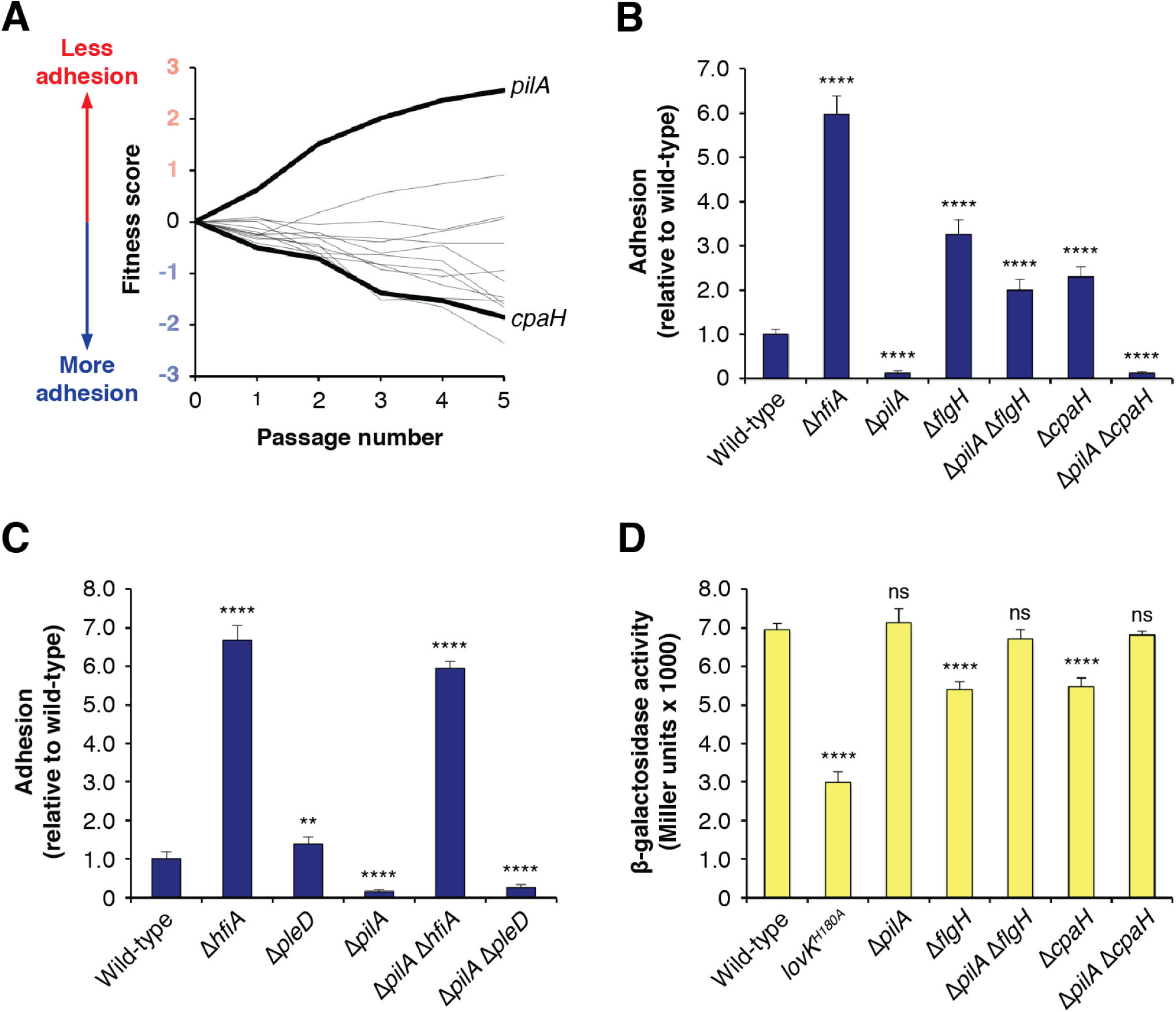
Opposing effects of pilus mutants on adhesion. A) Fitness profiles for genes at the pilus assembly locus. A full list of these genes and their annotations is provided in Table S5. B) Surface attachment of pilus mutants measured by CV staining. Cultures were grown for 17 hours in M2X medium before staining. Deletion of the gene for the main pilin subunit (PilA) reduces adhesion. Δ*pilA* is epistatic to Δ*cpaH* but not Δ*flgH.* The graph shows the average (± standard deviation) of seven biological replicates. C) Effect of *hfiA* and *pleD* deletions surface attachment in the Δ*pilA* background. Staining is slightly lower in Δ*hfiA* Δ*pilA* than Δ*hfiA* reflecting the holdfast-independent defect in surface attachment when the pilus is disrupted. *pleD* has no effect on adhesion in the Δ*pilA* mutant. The graph shows the average (± standard deviation) of six biological replicates. D) *P*_*hfiA*_*-lacZ* reporter activity in various *pilA* mutants. The chart shows the average (± standard deviation) of four biological replicates. Statistical significance was assessed by ANOVA with a pairwise Dunnett’s post test to determine which samples differed from wild type. ns – not significant; ** *P* < 0.01; **** *P* < 0.0001. When necessary, P values for additional pairwise comparisons pertinent to interpretation are indicated in the text.

To examine the relationship between *pilA-*dependent loss of adhesion and the activation of adhesion observed in mutants that disrupt pilus and flagellum assembly we created Δ*flgH* Δ*pilA* and Δ*cpaH* Δ*pilA* double mutants. The phenotypes for these mutants were similar in both complex and defined medium (Table S4). Surface attachment levels in the Δ*flgH* Δ*pilA* mutant were intermediate to those of Δ*pilA* (*P* < 0.0001) and Δ*flgH* (*P* < 0.0001), suggesting that *flgH* and *pilA* regulate adhesion through independent, additive pathways (Fig 4C and Table S4). In contrast, adhesion in the Δ*cpaH* Δ*pilA* mutant was indistinguishable from Δ*pilA* demonstrating that the effects of *pilA* on adhesion are epistatic to those of *cpaH* (Fig 4C and Table S4). We conclude that the pilin subunit PilA is required for the holdfast promoting effect caused by disruption of the pilus assembly apparatus.

We further explored the model that *pilA* contributed to the modulation of adhesion by *cpaH* by examining the relationships between *pilA, hfiA* and *pleD*. The decreased adhesion observed in the Δ*pilA* mutant was not affected by the subsequent deletion of *pleD,* indicating that the effect of *pilA* on adhesion is *pleD-*independent. Adhesion in a Δ*pilA* Δ*hfiA* double mutant was elevated to a level slightly below that of the Δ*hfiA* strain (*P* < 0.0001; Fig 4B). The difference in surface attachment between the Δ*pilA* Δ*hfiA* and Δ*hfiA* mutants likely reflects the holdfast-independent surface attachment defects caused by the loss of a functional pilus. Expression from *P*_*hfiA*_ was slightly elevated in the Δ*pilA* mutant (Fig 4D). However, because *hfiA* is already highly expressed under these conditions, it is difficult to determine if *P*_*hfiA*_ is activated further in the Δ*pilA* mutant. Finally, β-glactosidase activity from the *P*_*hfiA*_ reporter in both the Δ*flgH* Δ*pilA* and Δ*cpaH* Δ*pilA* backgrounds was restored to wild-type levels demonstrating that *pilA* is required to lower *hfiA* expression in the Δ*flgH* and Δ*cpaH* mutants (Fig 4D).

### New factors in the holdfast biosynthesis pathway

The final two clusters of mutants presented in Fig 1C displayed fitness profiles consistent with adhesion defects. We used the magnitude of the measured fitness changes to separate these genes into *1)* a cluster with higher fitness changes that contained all of the *hfs* genes known to be required for robust adhesion and *2)* a cluster displaying more modest fitness changes which contained the *hfsK* gene. *hfsK* encodes a putative *N*-acyltransferase thought to modify the holdfast polysaccharide in order to produce a fully adhesive holdfast (25). We chose three uncharacterized genes from these clusters of mutants to examine in detail for holdfast defects.

Disruption of *CCNA_01242,* which encodes a predicted amino acid permease, leads to the strongest non-adhesive fitness profile of any gene in the cheesecloth passaging experiment (Fig 5A). However, the Δ*CCNA_01242* strain had only a modest defect in surface attachment (Fig 5B). There were no obvious holdfast defects in the mutant, and we could not detect a significant adhesion defect under any conditions tested (Fig 5C and Table S2). Instead, Δ*CCNA_01242* had an unusual, biphasic growth profile. In complex medium, log phase was shorter than wild type, leading to a lower optical density as growth began to slow prematurely. Growth of this strain continued slowly over the next 24 h and eventually plateaued at a similar optical density to wild type (Fig S4). The biphasic growth seems to confound fitness calculations for samples collected during the sequential passaging experiment. It is not clear why *CCNA_01242* mutants were more enriched when cheesecloth was included in the medium, but we conclude nonetheless that this gene does not contribute to holdfast production.

**Figure 5.**
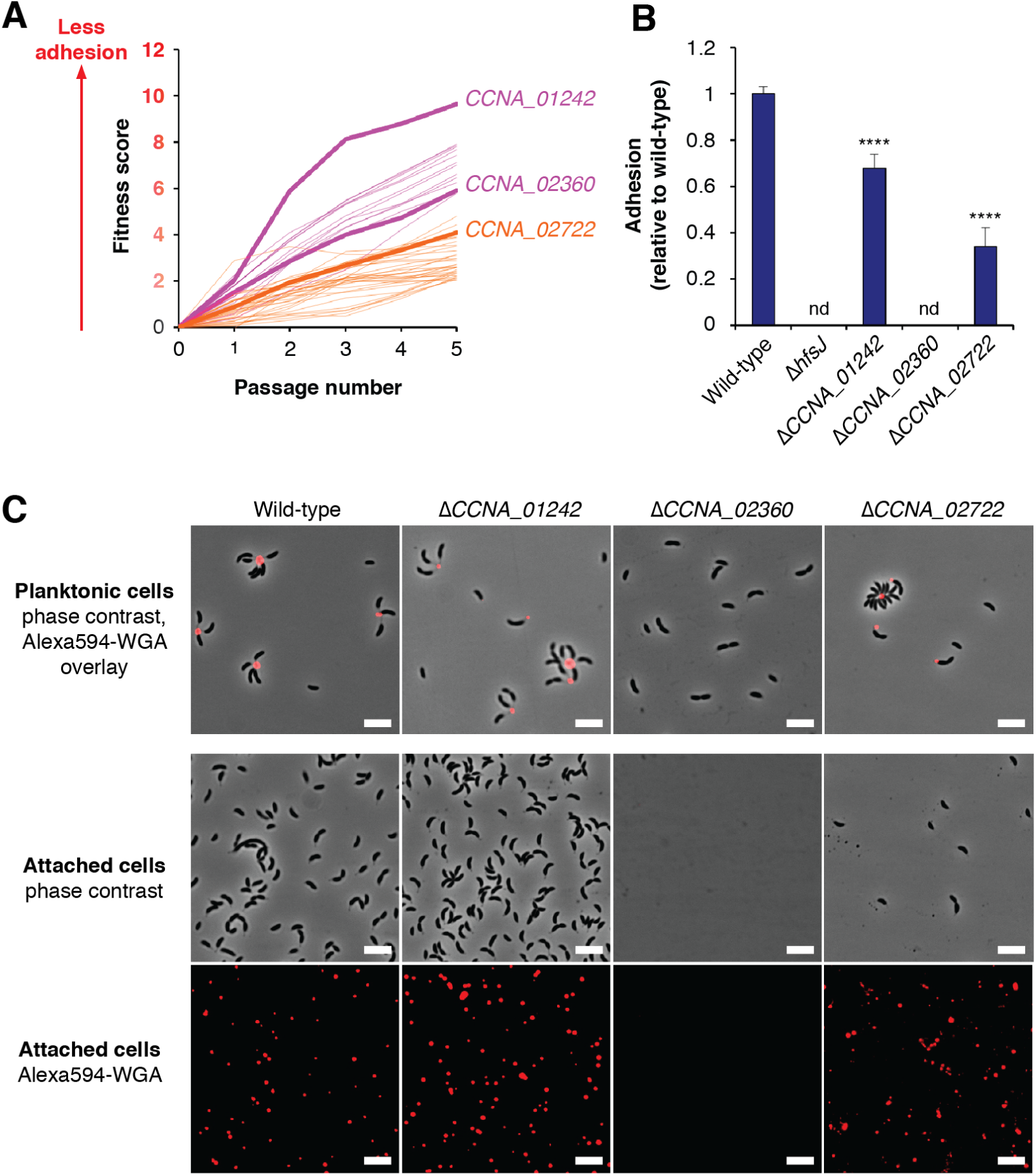
New holdfast biosynthesis factors. A) Fitness profiles for genes in the *hfs* (magenta) and holdfast modification (orange) clusters. Full lists of these genes and their annotations are provided in Table S5. B) Surface attachment of putative holdfast mutants measured by CV staining. Cultures were grown for 24 hours in PYE medium before being stained. Δ*CCNA_01242* and Δ*CCNA_02722* displayed reduced staining, and Δ*CCNA_02360* was non-adhesive. The graph shows the average (± standard deviation) of five biological replicates. Statistical significance was assessed by ANOVA with a pairwise Dunnett’s post test to determine which samples differed from wild type. nd – not detected; ns – not significant; **** *P* < 0.0001. C) Analysis of holdfast phenotypes by fWGA staining. The top panels show overlays of phase contrast and fluorescence images after staining planktonic cells as described in materials and methods. Adherent cells from the slide attachment assay are shown as phase contrast images in the middle set of panel, and the fluorescence channel showing attached holdfast material from the same slides is represented in the bottom panels. Δ*CCNA_01242* does not have an apparent holdfast defect, Δ*CCNA_02722* has a holdfast attachment defect and Δ*CCNA_02360* does not produce holdfasts. The scale bars represent 5μm.

*CCNA_02360* is a predicted member of the GT2 family of glycosyltransferases. Its disruptions have a fitness profile closely resembling many of the known *hfs* genes (Fig 5A). A Δ*CCNA_02360* mutant was non-adhesive in surface attachment assays and did not stain with fWGA under any conditions tested (Figs 5B and 5C). Given the lack of holdfast production in Δ*CCNA_02360* cells, we predict that *CCNA_02360* encodes a glycosyltransferase that contributes one or more monosaccharides to repeating unit of the holdfast polysaccharide and have named this gene *hfsL.* Previous studies have mentioned mutants in *CCNA_02360* as a holdfast-deficient control strain (25, 40), but, curiously, the identification of the gene and characterization of its phenotype have not been reported. Closely related genes could be identified in many stalked bacteria within the *Caulobacterales* clade suggesting that HfsL carries out a conserved step in holdfast biosynthesis. Identifying close homologs in more distantly related *Alphaproteobacteria* was difficult due to the abundance of GT2 family glycosyltransferases in bacterial genomes. Nevertheless, HfsL represents a fourth glycosyltransferase that is required for holdfast biosynthesis in *C. crescentus.*

*CCNA_02722* is a hypothetical protein that does not show homology to any known protein families. It has a predicted N-terminal signal peptide for export to the periplasm. The fitness profile is similar to *hfsK,* which is indicative of modest defects in adhesion (Fig 5A). The Δ*CCNA_02722* mutant was significantly impaired in surface-attachment (Fig 5B). When planktonic Δ*CCNA_02722* cells were stained with fWGA, holdfast staining was apparent (Fig 5C). However, using adhesion assays in which cells are grown in the presence of a glass slide that is then washed, stained with fWGA and imaged we detected a holdfast anchoring defect. Wild-type cells normally coat the surface of the slide and show fWGA foci at the site of attachment, but we observed very few attached Δ*CCNA_02722* cells. Instead, the slide was coated with an abundance of fWGA reactive material (Fig 5C). This holdfast shedding phenotype is characteristic of mutants with defects in anchoring the holdfast matrix to the surface of the cell (28), and the gene has been named *hfaE* accordingly. Like *hfsL, hfaE* could be identified in the genomes of many other stalked bacteria suggesting a conserved role in holdfast anchoring.

## Discussion

Holdfast production in *C. crescentus* presents an attractive model system to interrogate the assembly of polysaccharides in bacteria. Lectin staining, enzymatic sensitivity and functional annotations for the *hfs* genes indicate that the holdfast contains a polysaccharide (17, 18, 23). However, despite decades of work, surprisingly little else is known about the chemical structure of the holdfast. As part of our efforts to characterize the biosynthetic pathway, we sought a complete list of enzymes required for holdfast biosynthesis. In order to saturate the search for *hfs* genes, we developed a TnSeq-based method to measure the adhesion phenotype conferred by each non-essential gene in the genome.

The genome-wide approach provided a surprisingly rich set of insights into adhesion in *C. crescentus.* Not only did the screen identify non-adhesive mutants representing missing components of the holdfast pathway, but it also resolved mutants that displayed increased adhesion. One class of such mutants was defective in the production of smooth-LPS. Disrupting SLPS resulted in elevated adhesion levels in both holdfast producing and holdfast deficient backgrounds. SLPS mutants no longer adhered exclusively at the stalks of holdfast producing cells, but rather displayed a generalized form of adhesion throughout the cell surface. Our analysis of these mutants supports a model in which the *C. crescentus* envelope is structured to ensure that the cell surface is non-adhesive, maximizing the opportunity for polar adhesion via the holdfast. It is still unclear why disrupting the cell surface leads to the depletion and recovery profile seen during cheesecloth passaging. Regardless of the mechanism, simply identifying mutants that shared this temporal profile allowed for the characterization of new SLPS biosynthesis genes. Such co-fitness approaches have been useful in other contexts (42) and allowed us to greatly expand the number of genes in the *C. crescentus* SLPS pathway (Table S5).

Genes with predicted functions in motility, flagellar biosynthesis and type IV pilus assembly displayed a hyper-adhesive profile that could be distinguished from SLPS mutants by the lack of a recovery phase. Mutations in components of the flagellar basal body were recently shown to enhance holdfast production by inhibiting the expression of *hfiA,* a result that we confirmed here in our examination of *flgH* (41). Co-fitness analysis indicated that the *cpa* genes, which code for components of the type IV pilus, have a similar phenotype, and we showed that mutating the inner-membrane pilus assembly component *cpaH* also increased holdfast production by repressing *hfiA*. However, mutating *pilA,* which codes for the main subunit of the pilus filament, reduced adhesion. The adhesion defect in Δ*pilA* cells was partially restored in a Δ*pilA* Δ*flgH* background but remained unchanged in Δ*pilA* Δ*cpaH* cells. Thus, although the Δ*flgH* and Δ*cpaH* mutations both enhance holdfast production, the two pathways can be distinguished by their requirement for *pilA*. Disentangling the specific routes by which the various polar appendage mutants modulate *hfiA* activity will require identifying intermediate factors in the signaling pathways, but our results underscore the interconnectedness of the flagellum, pilus and holdfast.

Numerous reports have debated the roles of pili and the flagellum in surface attachment (13-15, 43, 44). Our unbiased, genome-wide analysis of adhesion unambiguously identified both appendages as determinants of attachment. Two recent studies, in particular, showed that mutating the flagellar basal body represses *hfiA* and that disruption of flagellar rotation upon surface contact stimulates holdfast production (15, 41). In our cheesecloth passaging experiment flagellar motor (*mot*), flagellin glycosylation (*flm)* and chemotaxis (*che*) genes shared the same hyper-adhesive fitness profile as components of the flagellar basal body (Table S5). Some of these mutants would be predicted to disrupt flagellar rotation without affecting assembly *per se* (45). We propose that disrupting flagellar function, either through physical interaction with a surface or mutation of motility genes, stimulates holdfast production. It will be interesting to test this model by determining whether the repression of *hfiA* seen in the flagellar mutants is required to activate holdfast production after surface contact. We also note that future studies should take into account the finding that mutating *flgH* reduces biofilm formation in a holdfast-independent manner (Fig S3), suggesting that flagellar motility likely also promotes productive interactions with a surface that lead to permanent attachment.

Much like flagellar mutants, disrupting components of the type IV pilus also causes both increased holdfast production and holdfast-independent surface colonization defects. A recent report proposed that contact with a surface inhibits the retraction of PilA filaments leading to a stimulation of holdfast production (14). One might initially conclude that pilus assembly defects in the *cpa* mutants mimic the obstruction of pilus filament retraction. However, the situation is more complex because *pilA* was required for increased holdfast production in the Δ*cpaH* mutant. These findings can be reconciled in a model in which the disruption of filament oscillation caused either by surface contact or upon mutation of the *cpa* genes leads to increases in the pool of unassembled PilA that serve as a signal to stimulate holdfast production. A similar model was proposed for the regulation of biofilm formation by the *Agrobacterium tumefaciens* pilus (38). Furthermore, unassembled pilin subunits in *Pseudomonas aeruginosa* directly activate the sensor kinase PilS leading to the repression of *pilA* transcription (46). Although no clear PilS homologs are found in *C. crescentus,* this example demonstrates that the membrane-associated pilin pool can serve as an input that activates signaling cascades.

An important aspect of both the flagellar and pilus pathways for holdfast regulation is their partial dependence on *pleD. pleD* is a pleiotropic regulator of cell polarity that is required for flagellar ejection and stalk synthesis during the swarmer to stalked cell transition (47). It functions as a diguanylate cyclase that is activated by phosphorylation at distinct stages of the cell cycle to produce the bacterial second messenger cyclic di-GMP (cdG) (39, 48). Though *pleD* has been shown to regulate the timing of holdfast synthesis during the cell cycle, Δ*pleD* mutants do not have significant bulk adhesion defects (Fig 3C) (40). More broadly, placing *pleD* within a signaling cascade that regulates holdfast synthesis is confounded by the fact that both *pleD* and cdG contribute to numerous processes that intersect with holdfast synthesis including flagellar function, pilus assembly, stalk biogenesis, and cell cycle progression (39, 49). Importantly, it has been shown that the holdfast glycosyltransferase HfsJ, which is the target of inhibition by HfiA, is also directly activated by cdG (11, 15). Thus, the role of PleD in promoting holdfast synthesis upon disruption of the pilus or flagellum is likely two-fold. It activates HfsJ by producing cdG and relieves inhibition by lowering transcription of *hfiA* through an unknown mechanism. PleD-dependent increases in cdG concentration may account for the disparity between the large changes in adhesion and modest decreases in *hfiA* expression in the Δ*flgH* and Δ*cpaH* backgrounds (Fig 3).

Finally, the initial goal of this study was to saturate the search for holdfast production factors. We used the cheesecloth passages to define a temporal fitness profile that was shared by genes known to be required for holdfast biosynthesis and used this pattern to search for missing genes in the pathway. In addition to a new holdfast anchoring factor, *hfaE*, we identified a glycosyltransferase, *hfsL,* and showed that it is required for holdfast production. Due to the saturating nature of our experiment, we believe that HfsE (along with the redundant PssY and PssZ), HfsJ, HfsG and HfsL carry out the only glycosyltransferase steps required for holdfast biosynthesis (Fig 6). Thus, we predict that a repeating unit of four or more monosaccharides makes up the holdfast polysaccharide because each of these enzymes likely contributes at least one sugar to the glycolipid intermediate that serves as a substrate for polymerization.

**Figure 6.**
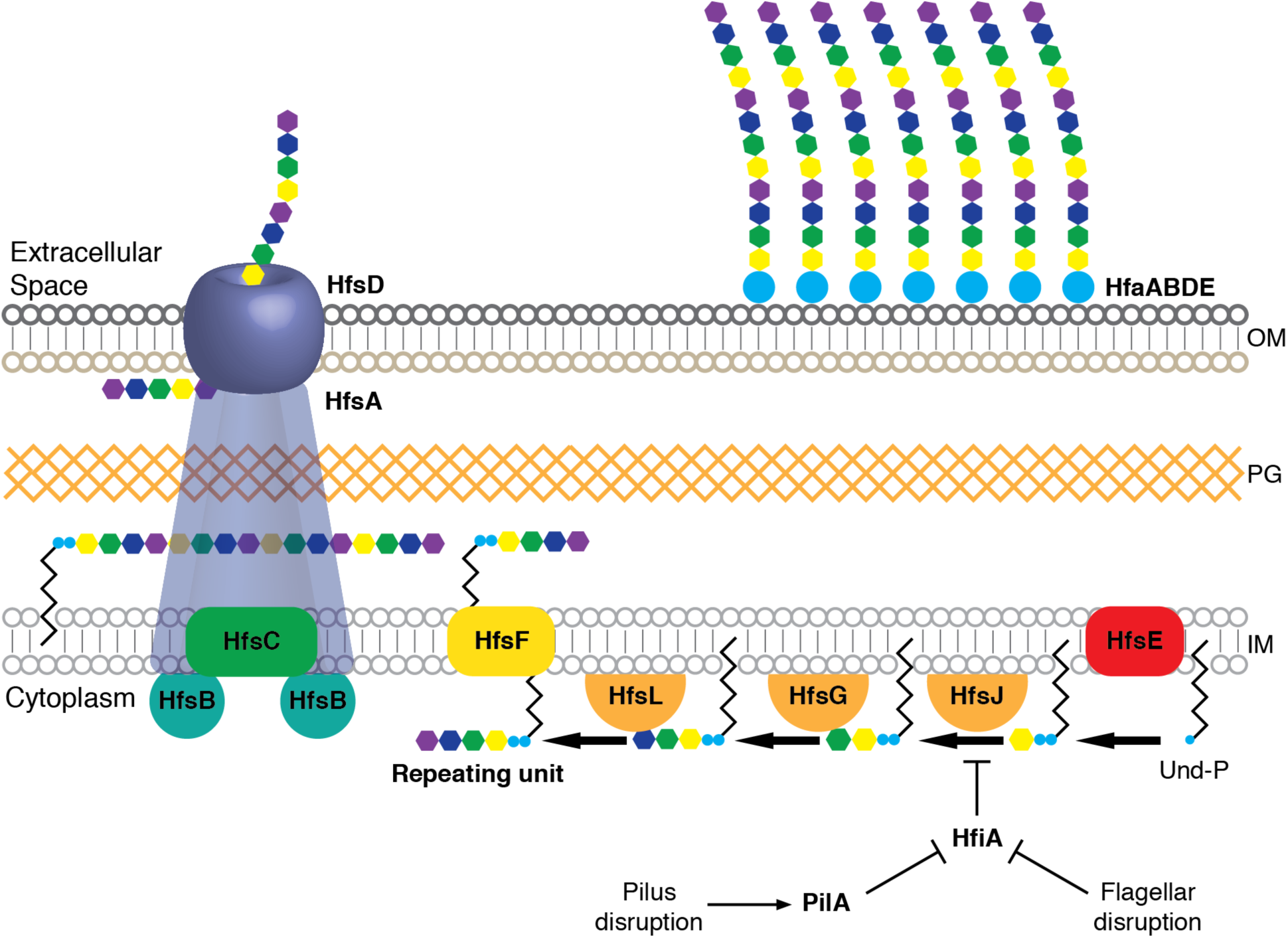
Updated model for holdfast biosynthesis. The model shows a *wzy-*type polysaccharide biosynthesis pathway. Four glycosyltransferases, HfsE, HfsJ, HfsG, and the newly described HfsL add monsaccharides sequentially onto the UP carrier to produce a glycolipid repeating unit. This intermediate is flipped across the membrane by HfsF, polymerized by HfsC and exported by a putative HfsABD transevelope complex. Attachment of the holdfast matrix is mediated by the Hfa proteins, including the newly-identified HfaE, reported here. Disruptions to the flagellum or the pilus activate holdfast production by relieving the inhibition of HfsJ by HfiA.

Implicit in defining the complete set of genes in the holdfast pathway is knowledge of genes that do not contribute. Many bacterial polysaccharides contain specialized monosaccharide components (50, 51), and these intermediates are often synthesized as nucleotide activated precursors that are directly utilized by glycosyltransferases (52, 53). We did not identify any genes involved in nucleotide sugar metabolism that had significant adhesion defects when disrupted. The modification factors HfsH and HfsK do likely convert certain monosaccharide components into more specialized residues. However, functional annotations for these enzymes predict that they act on lipid linked or polymerized substrates downstream of the glycosyltransferases, and neither enzyme is explicitly required for holdfast synthesis (19, 25). Our results suggest that specialized sugar precursors are not needed to produce the holdfast and that it is instead built using “housekeeping” sugars that are shared by other cellular processes. The ability to utilize standard nucleotide sugars as substrates without requiring specialized chemical syntheses will make the holdfast biosynthetic enzymes useful models for probing the catalytic mechanisms of bacterial polysaccharide biosynthesis.

The findings reported here highlight the advantages to probing mutant phenotypes in parallel. Classical genetic selections in which mutants are enriched then isolated and analyzed separately inherently favor the most extreme phenotypes. The single strain approach also favors longer genes that contribute more individual mutants to the pool. Screening mutants in parallel using TnSeq reduces these biases allowing detection of a range of phenotypes and capturing phenotypes for less abundant mutants in the pool. In this study, we discovered new regulatory networks that modulate holdfast synthesis by identifying mutants that display modest adhesion increases under conditions in which cells are already adhesive. Such mutants would be extremely difficult to isolate using a classical approach. Additionally, the saturating nature of these experiments makes them ideal for outlining complete biosynthetic pathways. Having a reasonable measure of saturation allowed us to leverage the results of the genome-wide screen to propose a model for the enzymatic steps in the holdfast pathway that will inform efforts to reconstitute the biosynthetic reactions *in vitro* (Fig 6).

## Methods

### Strains, growth conditions and genetic manipulation

Strains and plasmids used in this study are listed in Table S6. *E. coli* strains were grown in LB at 37°C with diaminopimelic acid (60 mM), kanamycin (50 μg/mL) or tetracycline (12 μg/mL) included as needed. *C. crescentus* CB15 was grown either in PYE (complex) or M2 salts containing 0.15% xylose (M2X, defined) at 30°C (54). When necessary sucrose (3%), kanamycin (25 μg/mL) or tetracycline (2 μg/mL) was included in solid medium. Kanamycin (5 μg/mL) or tetracycline (1 μg/mL) was included in liquid medium when necessary. Standard techniques were used for Gibson assembly-based cloning and sequencing plasmids (55). Primer sequences used for cloning specific constructs are available on request. Plasmids were introduced by electroporation or, in the case of the pFC1948 reporter plasmid, by tri-parental mating. Unmarked mutations were created through two-step deletion using a SacB counter-selection. Mutants were complemented by inserting target genes into pXGFPC-2, which allows for the integration of the plasmid at the *xyl* locus and transcriptional control from *P*_*xyl*_ (56). When necessary, target genes were inserted into pXGFPC-2 in reverse orientation under the control of their native promoters.

### Library development and mapping

The barcoded HiMar transposon pool APA_752 developed by Wetmore et al. (30) was used to create a barcoded Tn library in *C. crescentus* CB15. Construction of the library has been reported previously along with its associated statistics (57). Briefly, the transposon pool was introduced into *C. crescentus* by conjugation. Transconjugants appearing on selective plates were pooled, used to inoculate a liquid culture with Kanamycin and grown for 3 doublings. Glycerol was added to a final concentration of 15% and 1mL aliquots were frozen and stored at -80°C. TnSeq also followed the method of Wetmore et al (30). A 1mL library aliquot was centrifuged and genomic DNA was extracted from the pellet. The DNA was sheared, size selected for ∼300bp fragments and end repaired. A custom Y-adapter (Mod2_TS_Univ annealed to Mod2_TruSeq) was ligated and transposon junctions were amplified by PCR using the Nspacer_BarSeq_pHIMAR and P7_mod_TS_index1 primers. An Illumina HiSeq2500 was used to generate 150 bp single-end reads of the library. The genomic positions of each barcoded insertion were determined with BLAT. The barcode corresponding to each insertion site was determined using MapTnSeq.pl. This information was used to develop a list of barcodes that mapped to unique insertion sites using DesignRandomPool.pl (available at https://bitbucket.org/berkeleylab/feba).

### Passaging in cheesecloth

1 mL aliquots of the transposon library were thawed at room temperature and 300 μL of the library was inoculated into a well of a 12-well microtiter plate containing 1.2 mL PYE and a stack of 5 squares (∼10 mm x 10 mm) of sterile cheesecloth was added. The culture was grown with 155 rpm shaking at 30°C for 24 h after which 100 μL of the planktonic culture was used to inoculate a fresh well containing 1.4 mL PYE and a fresh piece of sterile cheesecloth. An additional 500 μL of the planktonic culture medium was centrifuged and the pellet was stored at -20°C for BarSeq analysis. The process was repeated for a total of five passages and each passaging experiment was performed in triplicate. The same procedure was used to perform passaging experiments in which no cheesecloth was added.

### Fitness determination with BarSeq

Cell pellets from 0.5mL of planktonic culture medium that had been frozen after each passage were used as templates for PCR reactions that simultaneously amplified the barcode region of the transposon insertions while adding TruSeq indexed Illumina adaptors (30). The PCR products were purified, pooled and multiplexed on a single Illumina 4000 lane for sequencing. Fitness values for each gene were determined using the pipeline described by Wetmore et al (30). Barcodes for each read were mapped using MultiCodes.pl and correlated with their associated insertion positions using combineBarSeq.pl. This data was used to calculate fitness using FEBA.R. This analysis determines strain fitness as the log_2_ ratio of barcode counts in a sample to the barcode counts in the reference condition, for which we used the first passage of the library in PYE without cheesecloth. Gene fitness was calculated by determining the weighted average of the insertion mutants with a given gene, excluding the first and last 10% of the ORF. The scripts used for fitness determination can be found at https://bitbucket.org/berkeleylab/feba.

### Data availability

The raw sequence data for each BarSeq sample along with a table describing the barcode abundances and their genomic insertion sites has been uploaded to the NCBI Gene Expression Omnibus (GEO) at https://www.ncbi.nlm.nih.gov/geo/ under the accession: GSE119738.

### Analysis of fitness data

We focused on identifying genes with the highest absolute (positive or negative) fitness scores. For each passaging step (with and without cheesecloth), an average and a standard deviation for the three replicate samples were calculated. Genes for which the largest standard deviation across the ten conditions was greater than the largest absolute fitness score were eliminated from further analysis. Fitness scores determined for each passage without cheesecloth were subtracted from those of the corresponding passage with cheesecloth to normalize for growth defects. These normalized values were used to rank each gene according to its largest absolute fitness score at any stage of cheesecloth passaging. The top 250 genes were then sorted by hierarchical clustering (58) to identify related fitness patterns. Groups derived from the clustering analysis were curated manually to produce the genes in Fig 1C. The “Pilus Assembly” subsection of Table S5 was created by manually identifying known pilus assembly genes, some of which do not have phenotypes that are strong enough to meet the cutoff used to identify the genes in Fig 1C.

### Crystal violet (CV) staining of adherent cells

Overnight cultures of *C. crescentus* grown in PYE were diluted to OD_660_=0.4 with PYE, 1uL was inoculated into the wells of 48-well microtiter plates containing 450uL medium and the cultures were shaken for 24 h. For cultures used to measure the effects of polar appendage mutants on adhesion, the incubation was shortened to 17 h. The cultures were then discarded, and the plates were washed thoroughly with a steady stream of tap water. Surface attached cells remaining in the wells were stained for 5 min with 0.01% crystal violet and washed again with tap water. Stain was extracted for 5 min in 100% EtOH and quantified by reading absorbance at 575nm.

### Fluorescent wheat germ agglutinin (fWGA) staining

For staining of planktonic cells, 400 μL of liquid culture was added to an Eppendorf tube and Alexa594 conjugated-WGA (ThermoFisher) was added to a final concentration 0.5 μg/mL. After incubating the cells for 5 min in the dark, 1mL of sterile water was added and the cells were centrifuged at 6k x g for 2 min. The pellet was re-suspended in 5 μL of medium, spotted on a glass slide and imaged. Cells were imaged with a Leica DM500 microscope equipped with a HCX PL APO 63X/ 1.4na Ph3 objective. fWGA staining was visualized using an RFP fluorescence filter (Chroma set 41043). For quantifying the number of holdfast-producing cells, a culture was inoculated to an OD_660_ of 0.002 and harvested when the OD_660_ reached 0.05-0.1 to minimize the number of rosettes.

For staining attached cells, a glass coverslip that had been washed with ethanol was added to 1mL of PYE in a 12-well plate, and 100uL of a starter culture (diluted to OD_660_=0.4) was added to the well. The cultures were grown for 6-8 h. Un-attached cells were removed from the coverslip by washing under a stream of distilled water. One side of the glass was covered with a solution of 0.5 μg/mL fWGA in PYE, incubated for 5 min in the dark and washed under a stream of distilled water. The coverslip was placed stain side down on a glass slide and imaged with phase contrast and fluorescence as described above.

### Smooth lipopolysaccharide (SLPS) Immunoblotting

Lysates for immunoblotting were prepared by the method of Walker et al (59). Pellets from saturated cultures were treated with DNAse and lysozyme, mixed with SDS-running buffer and digested with proteinase K. Samples were separated on a Tris-glycine SDS-PAGE gel with 12% acrylamide and transferred to nitrocellulose. The amount of sample loaded was normalized to the final optical density of the culture at harvest. The membrane was blocked in 5% milk, probed with a 1 in 20,000 dilution of anti-SLPS serum raised in rabbit (34), washed, probed with HRP-conjugated goat-anti-rabbit antibody (Invitrogen), washed again and visualized with peroxidase substrate. The top half of the membrane was removed before blocking and stained with Ponceau S (0.25% (w/v) Ponceau S in 1% acetic acid) as a loading control.

### Analysis of Rough LPS

LPS was extracted by the method of Darveau and Hancock (60). Cells were isolated from saturated 50mL *C. crescentus* cultures grown in M2X by centrifugation, re-suspended in 2 mL of 10 mM Tris-HCl pH 8.0 containing 2 mM MgCl_2_, and sonicated. DNAseI and RNAseA were added to 100 μg/mL and 25 μg/mL, respectively and the lysate was incubated for 1 hour at 37°C. Additional DNAseI and RNAseA were added to 200 μg/mL and 50 μg/mL, respectively and the lysate was incubated for an additional 1 hour at 37°C. SDS and EDTA were added to 2% and 100 mM, respectively, and the lysate was incubated for 2 h at 37°C. The solution was then centrifuged for 30 min at 50,000 x *g*. Proteinase K was added to 50 μg/mL in the supernatant and the solution was incubated for 2 h at 60°C after which the LPS was precipitated with 2 volumes of ice-cold 0.375 M MgCl_2_ in 95% EtOH and collected by centrifugation at 12,000 x *g*. The precipitate was re-suspended in 2% SDS containing 100mM EDTA, incubated overnight at 37°C, and re-precipitated with 0.375 M MgCl_2_ in 95% EtOH. The precipitate was then suspended in 10 mM Tris-HCl pH 8.0 and centrifuged for 2 h at 200,000 x *g*. The LPS pellets from each strain were suspended in SDS loading dye and separated on by Tris-Tricine SDS-PAGE on an 18% acrylamide gel containing 6M urea. LPS was stained using the periodate-silver method of Kittelberger and Hilbink (61).

### Bacteriophage FCBK sensitivity

Saturated cultures of *C. crescentus* grown in PYE were diluted to OD_660_=0.4 with PYE. 4μL of these suspensions along with six 10-fold serial dilutions (prepared in PYE) were spotted on PYE plates containing 0.15% xylose that had been spread to 107 PFU/mL (assuming a volume of 60mL for each 150mm plate) with ΦCBK or PYE plates with 0.15% xylose alone. The plates were incubated for 48 h at 30°C and photographed.

### Soft-agar swarming assay

1.5μL from a saturated culture of the appropriate *C. crescentus* strain grown in PYE was spotted in plates of PYE containing 0.3% agar and 0.15% xylose. The plates were incubated for 4 days at 30°C and photographed.

### lacZ reporter assay

Cultures for measuring reporter activity were grown in M2X medium and the amount of culture required to achieve an OD_660_ of 0.0005-0.00075 was added to fresh M2X medium. These cultures were grown to an OD_660_ of 0.05-0.15 and β-galactosidase activity was measured as previously described (11, 62).

## Supporting information

## Acknowledgements

We thank members of the Crosson lab and Prof. Arash Komeili for helpful discussions and insight. We thank John Smit for the gift of the SLPS anti-serum. This work was supported by NIH grant R01GM087353 to S.C. D.M.H. is supported by the Helen Hay Whitney Foundation.

