## Supplementary material for "A genome-wide analysis of adhesion in *Caulobacter crescentus* identifies new regulatory and biosynthetic components for holdfast assembly"

**Table S1** *Samples used for BarSeq analysis of gene fitness across cheesecloth passages*

The “Index” column represents the TruSeq indices used to de-multiplex the samples after sequencing. Cheese and PYE represent passages with and without cheesecloth, respectively. The numbers indicate which of the five passages each sample represents, and the letters indicate which of the three replicates. The “Proportion of reads in top percentile” column is calculated by ranking the barcodes in each sample by their abundance and determining the proportion of reads that map to the top 1% of barcodes.

| **Sample name** | **Index** | **Sample description** | **Total reads** | **Number of analyzable reads** | **Unique barcodes identified** | **Proportion of reads in top percentile** |
| --- | --- | --- | --- | --- | --- | --- |
| SCD_50.31 | CACGAT | PYE 1A | 4018890 | 3698106 | 233280 | 0.117 |
| SCD_50.32 | CACTCA | PYE 1B | 4492543 | 4130359 | 246118 | 0.1227 |
| SCD_50.33 | CAGGCG | PYE 1C | 4449306 | 4044087 | 150780 | 0.1236 |
| SCD_50.34 | CATGGC | PYE 2A | 4137845 | 3779486 | 226993 | 0.1249 |
| SCD_50.35 | CATTTT | PYE 2B | 4499135 | 4137664 | 228359 | 0.1274 |
| SCD_50.36 | CCAACA | PYE 2C | 4213003 | 3750243 | 228855 | 0.1316 |
| SCD_50.37 | CGGAAT | PYE 3A | 6735128 | 6171057 | 262618 | 0.1504 |
| SCD_50.38 | CTAGCT | PYE 3B | 6127591 | 5614129 | 249692 | 0.1471 |
| SCD_50.39 | CTATAC | PYE 3C | 5796738 | 5301288 | 246006 | 0.1563 |
| SCD_50.40 | CTCAGA | PYE 4A | 5322215 | 4857843 | 235579 | 0.1571 |
| SCD_50.41 | GACGAC | PYE 4B | 6394697 | 5855783 | 248801 | 0.1691 |
| SCD_50.42 | TAATCG | PYE 4C | 2975382 | 2755150 | 172812 | 0.1675 |
| SCD_50.43 | TACAGC | PYE 5A | 5799597 | 5301223 | 246151 | 0.2002 |
| SCD_50.44 | TATAAT | PYE 5B | 6016430 | 5569218 | 234947 | 0.1972 |
| SCD_50.45 | TCATTC | PYE 5C | 4811202 | 4428232 | 209890 | 0.2178 |
| SCD_50.46 | TCCCGA | Cheese 1A | 4562846 | 4108435 | 242916 | 0.1455 |
| SCD_50.47 | TCGAAG | Cheese 1B | 4313848 | 3922295 | 233599 | 0.1484 |
| SCD_50.48 | TCGGCA | Cheese 1C | 4773752 | 4335186 | 247032 | 0.1516 |
| SCD_50.49 | AAACAT | Cheese 2A | 6050903 | 5557666 | 232487 | 0.2488 |
| SCD_50.50 | AAAGCA | Cheese 2B | 7013967 | 6384659 | 253431 | 0.2793 |
| SCD_50.51 | AAATGC | Cheese 2C | 4421479 | 4054218 | 211068 | 0.2713 |
| SCD_50.52 | AACAAA | Cheese 3A | 6616953 | 5904400 | 235379 | 0.4118 |
| SCD_50.53 | AACTTG | Cheese 3B | 5935535 | 5450038 | 209707 | 0.4585 |
| SCD_50.54 | AAGACT | Cheese 3C | 7750956 | 7158882 | 204209 | 0.5887 |
| SCD_50.55 | AAGCGA | Cheese 4A | 4002747 | 3664336 | 167909 | 0.6349 |
| SCD_50.56 | AAGGAC | Cheese 4B | 3156990 | 2902069 | 151371 | 0.605 |
| SCD_50.57 | AATAGG | Cheese 4C | 1882499 | 1730719 | 115437 | 0.6777 |
| SCD_50.58 | ACAAAC | Cheese 5A | 2475323 | 2197424 | 99108 | 0.787 |
| SCD_50.59 | ACATCT | Cheese 5B | 2419554 | 2241238 | 99833 | 0.7796 |
| SCD_5060 | ACCCAG | Cheese 5C | 2865211 | 2583906 | 107021 | 0.8014 |

**Table S2** *Complementation of adhesion defects*

Normalized crystal violet staining values are shown as the average ± standard deviation from at least 4 biological replicates. All values shown reflect trends that were consistent across at least five independent experiments. Cells were grown for 17 hours in M2X or 24 hours in PYE medium before staining. n.m. – not measured.

| **Genotype** | **Deletion (PYE)** | **Empty vector (PYE)** | **Complement (PYE)** | **Deletion (M2X)** | **Empty Vector (M2X)** | **Complement (M2X)** |
| --- | --- | --- | --- | --- | --- | --- |
| Wild-type | 1.00 ± 0.03 | n.m. | n.m. | 1.00 ± 0.06 | n.m. | n.m. |
| ∆*hfiA* | 1.28 ± 0.16 | n.m. | n.m. | 4.34 ± 0.23 | n.m. | n.m. |
| ∆*hfsJ* | 0.00 ± 0.00 | n.m. | n.m. | 0.00 ± 0.00 | n.m. | n.m. |
| ∆*CCNA_*  *01242* | 0.68 ± 0.06 | 0.98 ± 0.06 | 0.98 ± 0.13 | 0.81 ± 0.19 | 0.74 ± 0.09 | 1.09 ± 0.17 |
| ∆*hfaE* | 0.34 ± 0.08 | 0.32 ± 0.12 | 1.02 ± 0.08 | 0.06 ± 0.03 | 0.05 ± 0.02 | 1.08 ± 0.19 |
| ∆*hfsL* | 0.00 ± 0.00 | 0.00 ± 0.00 | 1.00 ± 0.12 | 0.00 ± 0.01 | 0.00 ± 0.01 | 1.10 ± 0.21 |
| ∆*CCNA_*  *00497* | 1.12 ± 0.07 | 1.02 ± 0.07 | 1.08 ± 0.04 | 2.10 ± 0.23 | 3.47 ± 0.35 | 0.84 ± 0.10 |
| ∆*CCNA_*  *02386* | 0.72 ± 0.07 | 0.68 ± 0.08 | 1.10 ± 0.14 | 1.97 ± 0.30 | 2.73 ± 0.19 | 0.72 ± 0.13 |
| ∆*rfbB* | 1.34 ± 0.06 | 1.40 ± 0.03 | 1.08 ± 0.07 | 3.24 ± 0.71 | 2.14 ± 0.24 | 1.48 ± 0.31 |
| ∆*wbqP* | 1.29 ± 0.11 | 1.09 ± 0.11 | 1.11 ± 0.12 | 3.87 ± 0.51 | 2.31 ± 0.40 | 1.10 ± 0.26 |
| ∆*flgH* | 0.58 ± 0.05 | 0.62 ± 0.06 | 1.03 ± 0.04 | 2.91 ± 0.08 | 2.42 ± 0.26 | 1.23 ± 0.25 |
| ∆*cpaH* | 0.71 ± 0.05 | 0.76 ± 0.08 | 1.03 ± 0.07 | 2.68 ± 0.07 | 1.85 ± 0.24 | 0.90 ± 0.31 |
| ∆*pilA* | 0.34 ± 0.04 | 0.44 ± 0.04 | 1.01 ± 0.06 | 0.20 ± 0.16 | 0.08 ± 0.05 | 0.93 ± 0.36 |

**Table S3** *Holdfast counts for polar appendage mutants­­*

The values represent the fraction of cells (out of 1) that stained with a holdfast focus along with the associated standard deviation from three biological replicates. The total number of cells counted is shown in parentheses. Cells were harvested from low-density cultures (Materials and Methods) to minimize the formation of rosettes. However, in the event that rosettes were observed, all cells in the rosette were counted as holdfast producing.

| **Strain** | **PYE** | **M2X** |
| --- | --- | --- |
| Wild-type | 0.686 ± 0.045 (739) | 0.081 ± 0.006 (743) |
| ∆*hfiA* | 0.863 ± 0.021 (526) | 0.722 ± 0.039 (703) |
| ∆*hfsJ* | 0.000 ± 0.000 (628) | 0.000 ± 0.000 (638) |
| ∆*flgH* | 0.869 ± 0.037 (662) | 0.620 ± 0.013 (609) |
| ∆*cpaH* | 0.874 ± 0.031 (531) | 0.296 ± 0.094 (689) |
| ∆*pilA* | 0.673 ± 0.030 (464) | 0.092 ± 0.016 (367) |
| ∆*pleD* | 0.496 ± 0.024 (457) | 0.084 ± 0.007 (500) |

**Table S4** *Additional adhesion phenotypes for polar appendage mutants*

Normalized crystal violet staining values are shown as the average ± standard deviation from at least 4 biological replicates. All values shown reflect trends that were consistent across at least five independent experiments. Cells were grown for 17 hours in M2X or 24 hours in PYE medium before staining. n.m. – not measured.

| **Genotype** | **Deletion (PYE)** | **∆hfiA (PYE)** | **∆pleD (PYE)** | **Deletion (M2X)** | **∆hfiA (M2X)** | **∆pleD (M2X)** |
| --- | --- | --- | --- | --- | --- | --- |
| Wild-type | 1.00 ± 0.03 | 1.28 ± 0.16 | 0.81 ± 0.01 | 1.00 ± 0.06 | 5.61 ± 0.27 | 1.38 ± 0.19 |
| ∆*hfiA* | 1.28 ± 0.16 | n.m. | 1.08 ± 0.07 | 4.34 ± 0.23 | n.m. | 5.53 ± 0.16 |
| ∆*hfsJ* | 0.00 ± 0.00 | n.m. | n.m. | 0.00 ± 0.00 | n.m. | n.m. |
| ∆*wbqP* ∆*hfsJ* | 0.48 ± 0.25 | n.m. | n.m. | 2.51 ± 0.64 | n.m. | n.m. |
| ∆*flgH* | 0.58 ± 0.05 | 0.62 ± 0.07 | 0.31 ± 0.02 | 2.91 ± 0.08 | 2.78 ± 0.21 | 1.41 ± 0.33 |
| ∆*cpaH* | 0.71 ± 0.05 | 0.91 ± 0.05 | 0.19 ± 0.01 | 2.68 ± 0.07 | 4.29 ± 0.16 | 1.22 ± 0.18 |
| ∆*pilA* | 0.34 ± 0.04 | 0.73 ± 0.03 | 0.43 ± 0.01 | 0.20 ± 0.16 | 5.95 ± 0.18 | 0.25 ± 0.08 |
| ∆*flgH* ∆*hfsJ* | 0.00 ± 0.00 | n.m. | n.m. | 0.01 ± 0.01 | n.m. | n.m. |
| ∆*cpaH* ∆*hfsJ* | 0.00 ± 0.00 | n.m. | n.m. | 0.01 ± 0.02 | n.m. | n.m. |
| ∆*flgH* ∆*cpaH* | 0.40 ± 0.01 | n.m. | n.m. | 4.13 ± 0.18 | n.m. | n.m. |
| ∆*flgH* ∆*pilA* | 0.32 ± 0.02 | n.m. | n.m. | 2.00 ± 0.25 | n.m. | n.m. |
| ∆*cpaH* ∆*pilA* | 0.43 ± 0.03 | n.m. | n.m. | 0.12 ± 0.04 | n.m. | n.m. |

**Table S5** *Fitness scores across cheesecloth passages for mutant clusters shown in Fig 1C*

Fitness values represent the average of the three replicates for each passage. The first passage without cheesecloth (PYE1) represents the time 0 sample.

| ***Cluster 1: SLPS*** |  |  |  |  |  |  |
| --- | --- | --- | --- | --- | --- | --- |
| **locusId; annotation** | **0** | **1** | **2** | **3** | **4** | **5** |
| CCNA_00044; ribosome maturation protein RimP | 0.013 | -2.93 | -4.145 | -3.319 | -1.57 | -0.067 |
| CCNA_00217; thiol:disulfide interchange protein DsbD | -0.07 | -2.488 | -2.869 | -1.883 | -0.313 | 1.215 |
| CCNA_00290; autotransporter protein | 0.009 | -3.141 | -3.74 | -3.294 | -2.788 | -1.794 |
| CCNA_00390; ADP-heptose--LPS heptosyltransferase | -0.008 | -2.321 | -4.402 | -2.913 | -2.413 | -1.408 |
| CCNA_00497; putative rhamnosyl transferase | -0.074 | -3.219 | -3.706 | -2.835 | -1.19 | 0.371 |
| CCNA_00502; glycosyl transferase family protein | 0.024 | -3.405 | -3.06 | -3.883 | -2.296 | -1.313 |
| CCNA_00512; GTP-binding protein, probable translation factor | 0.082 | -3.203 | -3.936 | -3.087 | -1.455 | 0.104 |
| CCNA_00519; conserved hypothetical protein | 0.009 | -4.406 | -4.912 | -4.798 | -3.134 | -1.521 |
| CCNA_00667; lipopolysaccharide biosynthesis protein | -0.194 | -2.497 | -2.358 | -1.846 | -0.306 | 1.286 |
| CCNA_00668; capsular polysaccharide biosynthesis protein | -0.001 | -2.031 | -3.171 | -3.547 | -1.836 | -1.285 |
| CCNA_00669; glycosyltransferase family 99 protein WbsX | -0.022 | -3.396 | -3.205 | -2.343 | -0.788 | 0.792 |
| CCNA_01055; GT1 family glyscosyl transferase | -0.043 | -3.088 | -4.106 | -3.578 | -2.06 | -0.533 |
| CCNA_01062; GDP-mannose 4,6 dehydratase | -0.117 | -3.685 | -4.072 | -3.613 | -1.995 | -0.595 |
| CCNA_01063; UDP-perosamine 4-acetyl transferase | -0.038 | -3.41 | -2.969 | -2.665 | -0.986 | 0.2 |
| CCNA_01064; perosamine synthetase | -0.112 | -3.481 | -4 | -3.214 | -1.594 | -0.064 |
| CCNA_01065; glycosyltransferase | -0.076 | -3.761 | -3.547 | -3.178 | -1.581 | -0.043 |
| CCNA_01066; glycosyltransferase | -0.038 | -2.802 | -3.277 | -2.57 | -1.632 | -0.394 |
| CCNA_01068; glycosyltransferase | -0.15 | -4.496 | -3.787 | -3.662 | -1.324 | -0.508 |
| CCNA_01086; GTP-binding protein lepA | 0.043 | -3.449 | -3.931 | -3.509 | -2.112 | -0.57 |
| CCNA_01103; ADP-heptose--LPS heptosyltransferase | 0.034 | -2.691 | -2.823 | -3.069 | -1.778 | -0.077 |
| CCNA_01199; glucose-1-phosphate thymidylyltransferase | -0.008 | -4.037 | -3.851 | -3.099 | -1.424 | 0.062 |
| CCNA_01375; lactoylglutathione lyase | 0.048 | -3.392 | -4.654 | -4.066 | -2.463 | -1.058 |
| CCNA_01427; beta-barrel assembly machine (BAM) protein BamE | 0.052 | -2.633 | -4.028 | -4.39 | -2.727 | -1.235 |
| CCNA_01430; conserved hypothetical protein | -0.013 | -1.923 | -2.959 | -3.144 | -2.08 | -0.903 |
| CCNA_01447; homoserine dehydrogenase | 0.02 | -2.348 | -3.724 | -2.91 | -1.286 | 0.261 |
| CCNA_01497; ADP-L-glycero-D-manno-heptose-6-epimerase | 0.022 | -2.19 | -2.593 | -2.199 | -1.98 | -0.42 |
| CCNA_01955; zinc metalloprotease | 0.005 | -2.151 | -2.316 | -2.232 | -1.865 | -1.314 |
| CCNA_01971; peptidyl-prolyl cis-trans isomerase | 0.013 | -2.17 | -2.493 | -1.791 | -0.927 | 0.591 |
| CCNA_02219; hypothetical protein | 0.017 | -2.85 | -2.949 | -2.521 | -2.268 | -1.687 |
| CCNA_02326; acetylornithine aminotransferase/succinyldiaminopimelate aminotransferase | 0.019 | -1.712 | -2.166 | -1.194 | -0.525 | 0.791 |
| CCNA_02347; phosphomannomutase/phosphoglucomutase | -0.064 | -4.09 | -4.85 | -4.033 | -2.455 | -0.889 |
| CCNA_02386; O-antigen ligase related enzyme | -0.134 | -2.762 | -3.457 | -2.585 | -1.025 | 0.537 |
| CCNA_02463; UDP-N-acetylglucosamine 4-epimerase | 0.029 | -2.627 | -3.781 | -3.758 | -2.374 | -0.882 |
| CCNA_02650; N-acetyl-anhydromuramyl-L-alanine amidase | -0.018 | -1.617 | -3.035 | -2.194 | -0.678 | 0.911 |
| CCNA_02941; transcription elongation factor greA | -0.119 | -1.233 | -2.748 | -2.498 | -0.92 | 0.629 |
| CCNA_03026; two-component response regulator petR | -0.055 | -2.619 | -4.063 | -3.652 | -2.577 | -1.705 |
| CCNA_03195; RNA polymerase sigma factor RpoH | -0.001 | -2.089 | -3.735 | -3.307 | -1.951 | -1.294 |
| CCNA_03352; YebC/PmpR transcriptional regulator | -0.009 | -3.942 | -4.169 | -3.705 | -2.805 | -1.706 |
| CCNA_03475; homoserine kinase | -0.028 | -1.569 | -3.168 | -2.323 | -0.697 | 0.84 |
| CCNA_03609; outer membrane protein | -0.011 | -2.28 | -3.936 | -3.203 | -1.489 | -0.261 |
| CCNA_03705; conserved hypothetical protein | 0.053 | -1.304 | -3.383 | -1.826 | -0.864 | 0.665 |
| CCNA_03713; RNA polymerase sigma-54 factor rpoN | 0.02 | -1.566 | -3.418 | -2.288 | -1.299 | 0.198 |
| CCNA_03733; mannose-1-phosphate guanylyltransferase | -0.137 | -2.754 | -3.111 | -2.202 | -1.085 | 0.29 |
| CCNA_03744; dTDP-glucose 4,6-dehydratase | -0.009 | -2.964 | -3.124 | -1.88 | -1.002 | 0.388 |
| CCNA_03748; dTDP-4-dehydrorhamnose 3,5-epimerase | -0.059 | -2.155 | -2.242 | -1.967 | -0.249 | 0.731 |
| CCNA_03859; two-component response regulator cenR | 0.01 | -1.898 | -3.244 | -2.126 | -0.879 | 0.579 |
| CCNA_03909; conserved hypothetical protein | 0.028 | -2.113 | -4.213 | -2.917 | -1.893 | -0.234 |
| CCNA_03984; hypothetical protein | 0.007 | -1.779 | -2.747 | -2.011 | -0.292 | -0.697 |
| ***Cluster 2: Polar appendages*** |  |  |  |  |  |  |
| **locusId; annotation** | **0** | **1** | **2** | **3** | **4** | **5** |
| CCNA_00233; UDP-N-acetylglucosamine 4,6-dehydratase | 0.008 | -0.398 | -0.647 | -1.464 | -1.536 | -1.994 |
| CCNA_00234; WecE-family cell wall biogenesis enzyme | 0.005 | -0.587 | -1.119 | -1.518 | -1.539 | -2.71 |
| CCNA_00444; chemotaxis protein methyltransferase | -0.017 | -0.73 | -1.261 | -1.337 | -1.85 | -2.405 |
| CCNA_00447; chemotaxis protein cheD | -0.008 | -0.393 | -1.262 | -1.868 | -2.689 | -2.926 |
| CCNA_00449; cheYIII | -0.001 | -0.649 | -1.243 | -1.71 | -2.675 | -2.985 |
| CCNA_00542; hypothetical protein | 0.003 | -0.471 | -1.169 | -1.452 | -1.965 | -2.529 |
| CCNA_00787; chemotaxis motA protein | -0.012 | -0.721 | -0.806 | -1.894 | -2.588 | -3.396 |
| CCNA_00821; hypothetical protein | -0.027 | -0.613 | -1.472 | -0.837 | -1.245 | -2.046 |
| CCNA_00823; LuxR-like DNA-binding protein | -0.058 | -0.97 | -1.218 | -1.518 | -2.066 | -2.612 |
| CCNA_00942; flagellar hook-associated protein FlgL | 0.004 | -0.424 | -0.765 | -1.385 | -1.835 | -2.326 |
| CCNA_00943; flagellar hook-associated protein FlaN | -0.032 | 0.105 | -0.548 | -0.928 | -2.707 | -2.553 |
| CCNA_01004; flagellar basal-body rod protein FlgB | 0.004 | -0.37 | -0.935 | -1.357 | -2.617 | -3.031 |
| CCNA_01005; flagellar basal-body rod protein flgC | 0.004 | -0.426 | -0.738 | -1.599 | -2.121 | -3.296 |
| CCNA_01094; hypothetical protein | -0.002 | 0.146 | -0.421 | -1.277 | -1.473 | -3.97 |
| CCNA_01117; conserved hypothetical protein | 0.006 | -0.472 | -0.715 | -1.256 | -1.147 | -1.777 |
| CCNA_01524; FlbA protein | 0.001 | -0.427 | -0.921 | -1.919 | -1.998 | -2.886 |
| CCNA_01527; flagellin fljL | 0.011 | -0.601 | -1.477 | -1.912 | -2.791 | -2.989 |
| CCNA_01530; flagellin FljJ | -0.005 | -0.245 | -0.754 | -1.332 | -1.409 | -2.203 |
| CCNA_01532; regulatory protein flaY | 0.004 | -0.56 | -0.887 | -1.34 | -1.497 | -1.985 |
| CCNA_01562; 4-hydroxy-2-oxoglutarate aldolase/2-dehydro-3-deoxyphosphogluconate aldolase | -0.001 | 0.111 | -0.489 | -1.482 | -2.752 | -3.307 |
| CCNA_01644; chemotaxis motB protein | -0.002 | -0.479 | -0.629 | -1.493 | -2.599 | -2.458 |
| CCNA_01675; outer membrane protein | -0.008 | -0.328 | -0.695 | -1.409 | -1.926 | -2.472 |
| CCNA_01676; conserved hypothetical protein | 0.006 | -0.455 | -0.55 | -1.207 | -2.14 | -2.362 |
| CCNA_02142; flagellar basal-body rod protein flgF | -0.011 | -0.368 | -1.402 | -1.304 | -2.054 | -3.858 |
| CCNA_02143; flagellar basal-body rod protein flgG | -0.008 | -0.273 | -1.237 | -1.728 | -2.163 | -2.703 |
| CCNA_02144; flagella basal body P ring formation protein flgA | 0.004 | -0.357 | -0.887 | -1.701 | -1.911 | -3.179 |
| CCNA_02145; flagellar L-ring protein flgH | -0.005 | -0.473 | -1.193 | -1.891 | -2.175 | -3.406 |
| CCNA_02322; Co2+/Mg2+ efflux protein ApaG | 0.01 | -0.146 | -0.529 | -0.943 | -1.572 | -2.072 |
| CCNA_02411; putative lytic transglycosylase PleA | 0.008 | -0.039 | -0.349 | -0.84 | -1.079 | -1.064 |
| CCNA_02526; dihydroorotase | 0 | -0.85 | -0.636 | -1.913 | -2.573 | -3.177 |
| CCNA_02667; flagellar basal-body protein FlbY | -0.011 | -0.328 | -0.781 | -1.413 | -1.898 | -3.416 |
| CCNA_02796; conserved hypothetical protein | -0.019 | -0.623 | -1.316 | -2.031 | -2.836 | -4.51 |
| CCNA_02946; spsF-related cytidylyltransferase | 0.002 | -0.342 | -0.765 | -1.462 | -1.667 | -1.885 |
| CCNA_02947; spsG-related polysaccharide biosynthesis protein | 0.007 | -0.126 | -0.843 | -0.56 | -2.42 | -2.072 |
| CCNA_02950; hypothetical protein | 0.006 | -0.564 | -1.334 | -1.874 | -2.604 | -2.534 |
| CCNA_02951; WbqC-like family protein | 0.011 | -0.302 | -0.744 | -1.461 | -1.905 | -2.271 |
| CCNA_02961; NeuB-family N-acetylneuraminate synthase | 0.005 | -0.404 | -0.841 | -1.145 | -2.067 | -2.738 |
| CCNA_03035; TadC-related pilus assembly protein | 0.013 | -0.501 | -0.716 | -1.382 | -1.531 | -1.851 |
| CCNA_03036; TadB-related pilus assembly protein | 0.004 | -0.411 | -0.608 | -1.411 | -1.647 | -2.348 |
| CCNA_03041; pilus assembly protein CpaB | 0.004 | -0.193 | -0.661 | -0.824 | -0.946 | -1.653 |
| CCNA_03890; conserved hypothetical protein | 0.007 | 0.049 | -0.268 | -1.598 | -1.602 | -2.476 |
| ***Cluster 3: Pilus assembly*** |  |  |  |  |  |  |
| **locusId; annotation** | **0** | **1** | **2** | **3** | **4** | **5** |
| CCNA_03033; Flp pilus assembly protein TadD | 0.008 | 0.009 | -0.62 | -0.629 | -0.758 | -1.594 |
| CCNA_03035; TadC-related pilus assembly protein | 0.013 | -0.501 | -0.716 | -1.382 | -1.531 | -1.851 |
| CCNA_03036; TadB-related pilus assembly protein | 0.004 | -0.411 | -0.608 | -1.411 | -1.647 | -2.348 |
| CCNA_03037; pilus assembly ATPase CpaF | 0.005 | 0.095 | -0.253 | -0.32 | -0.409 | -0.403 |
| CCNA_03038; pilus assembly ATPase CpaE | -0.004 | -0.237 | -0.211 | -0.606 | -0.457 | -1.147 |
| CCNA_03039; pilus assembly protein CpaD | 0.037 | -0.125 | -0.317 | -0.922 | -1.06 | -0.943 |
| CCNA_03040; outer membrane pilus secretion channel CpaC | 0.008 | -0.244 | -0.478 | -0.827 | -1.23 | -1.472 |
| CCNA_03041; pilus assembly protein CpaB | 0.004 | -0.193 | -0.661 | -0.824 | -0.946 | -1.653 |
| CCNA_03042; pilus assembly prepilin peptidase CpaA | -0.001 | -0.267 | 0.188 | 0.554 | 0.736 | 0.911 |
| CCNA_03043; type IV pilin protein pilA | 0.006 | 0.615 | 1.513 | 2.023 | 2.366 | 2.569 |
| CCNA_03044; CpaC-related secretion pathway protein | 0.008 | -0.301 | -0.433 | -1.517 | -1.481 | -1.527 |
| CCNA_03045; TadG-related pilus assembly protein | 0 | 0.045 | -0.047 | 0.016 | -0.154 | 0.109 |
| CCNA_03046; TadE-related pilus assembly protein | -0.004 | -0.33 | -0.304 | -0.389 | -0.182 | 0.06 |
| ***Cluster 4: Holdfast synthesis*** |  |  |  |  |  |  |
| **locusId; annotation** | **0** | **1** | **2** | **3** | **4** | **5** |
| CCNA_00094; WecG/TagA-family glycosyltransferase HfsJ | -0.001 | 1.792 | 3.826 | 5.461 | 6.707 | 7.85 |
| CCNA_01241; Zn-dependent hydrolase, glyoxalase II family | -0.014 | 0.565 | 2.753 | 4.296 | 5.334 | 6.595 |
| CCNA_01242; amino acid permease | 0.008 | 2.034 | 5.865 | 8.127 | 8.789 | 9.649 |
| CCNA_02360; glycosyl transferase family 2 protein | 0.006 | 1.507 | 2.857 | 3.987 | 4.727 | 5.919 |
| CCNA_02436; hypothetical protein | -0.011 | 0.37 | 1.598 | 2.491 | 4.207 | 5.78 |
| CCNA_02509; glycosyltransferase hfsG | 0 | 1.44 | 2.961 | 4.334 | 5.264 | 6.452 |
| CCNA_02510; oligosaccharide deacetylase hfsH | -0.002 | 1.328 | 3.155 | 4.619 | 5.782 | 6.993 |
| CCNA_02512; polysaccharide autokinase-related protein hfsB | -0.005 | 1.819 | 3.721 | 5.31 | 6.57 | 7.798 |
| CCNA_02513; holdfast synthesis protein HfsA | -0.002 | 1.823 | 3.702 | 5.359 | 6.375 | 7.653 |
| CCNA_02514; polysaccharide secretin protein hfsD | 0.001 | 1.8 | 3.812 | 5.491 | 6.638 | 7.911 |
| CCNA_02567; sensory transduction histidine kinase pleC | 0.002 | 2.23 | 4.093 | 5.298 | 6.293 | 7.414 |
| CCNA_02711; holdfast attachment protein hfaA | -0.019 | 1.335 | 2.869 | 4.649 | 5.335 | 6.242 |
| CCNA_02712; holdfast attachment protein hfaB | 0.001 | 1.441 | 3.288 | 4.825 | 5.894 | 7.119 |
| CCNA_02713; holdfast attachment protein hfaD | 0.004 | 1.053 | 2.76 | 3.972 | 5.004 | 5.894 |
| ***Cluster 5: Holdfast modification*** |  |  |  |  |  |  |
| **locusId; annotation** | **0** | **1** | **2** | **3** | **4** | **5** |
| CCNA_00006; enoyl-CoA hydratase | 0.006 | 1.416 | 2.106 | 2.633 | 2.838 | 2.885 |
| CCNA_00011; chaperone protein DnaJ | -0.002 | 0.391 | 1.424 | 2.487 | 3.42 | 3.55 |
| CCNA_00134; surface protein | 0.012 | 1.004 | 1.223 | 2.175 | 2.703 | 3.104 |
| CCNA_00135; trypsin-like peptidase | 0.024 | 1.408 | 1.849 | 2.536 | 2.983 | 3.508 |
| CCNA_00247; two-component receiver protein SpdR | -0.022 | 0.746 | 1.113 | 2.001 | 1.936 | 2.122 |
| CCNA_00527; conserved hypothetical protein | 0.015 | 0.266 | 0.896 | 2.136 | 3.469 | 4.796 |
| CCNA_00543; methyl-accepting chemotaxis protein | 0.009 | 1.134 | 1.716 | 2.128 | 2.157 | 2.226 |
| CCNA_00551; hypothetical protein | -0.004 | 0.866 | 1.543 | 2.139 | 2.308 | 2.69 |
| CCNA_00554; methyltransferase | -0.001 | 0.062 | 0.822 | 2.006 | 2.545 | 2.978 |
| CCNA_00908; 3-oxoacyl-(acyl-carrier-protein) synthase III | 0 | 0.407 | 0.586 | 0.845 | 1.489 | 2.322 |
| CCNA_00948; CtrA inhibitory protein SciP | 0.007 | 1.162 | 1.949 | 2.067 | 2.008 | 2.039 |
| CCNA_01020; LacI-family transcriptional regulator | 0.007 | 0.551 | 1.386 | 2.301 | 2.185 | 2.402 |
| CCNA_01214; YjgP/YjgQ family membrane permease | 0.015 | 2.881 | 3.481 | 3.156 | 3.126 | 3.309 |
| CCNA_01215; histidine triad (HIT) hydrolase | 0.028 | 1.556 | 1.748 | 1.705 | 2.032 | 2.724 |
| CCNA_01345; short chain dehydrogenase | 0.005 | 0.092 | 0.3 | 0.598 | 1.441 | 2.418 |
| CCNA_01354; myo-inositol 2-dehydrogenase IdhA | 0.011 | 0.138 | 0.551 | 0.717 | 1.425 | 2.056 |
| CCNA_01893; SnoaL-like domain protein | -0.014 | 0.189 | 0.498 | 0.484 | 1.23 | 2.187 |
| CCNA_02087; deoxyguanosinetriphosphate triphosphohydrolase | 0.002 | 0.592 | 1.448 | 1.871 | 2.344 | 2.614 |
| CCNA_02125; polar development protein podJ | 0.008 | 1.149 | 2.18 | 2.796 | 3.456 | 4.419 |
| CCNA_02242; PHB granule-associated protein, phasin2 | -0.018 | 0.695 | 1.362 | 1.519 | 1.819 | 2.214 |
| CCNA_02415; Xre-family transcriptional regulator | 0.001 | -0.156 | 2.237 | 3.038 | 2.641 | 2.593 |
| CCNA_02619; stomatin/prohibitin-related protein | -0.013 | 0.45 | 1.896 | 3.271 | 3.38 | 3.866 |
| CCNA_02722; conserved hypothetical protein | 0.002 | 0.867 | 1.936 | 2.676 | 3.336 | 4.092 |
| CCNA_02846; DegP/HtrA-family serine protease | 0.018 | 0.267 | 1.041 | 1.554 | 1.965 | 2.587 |
| CCNA_02880; terminase-like family protein | -0.003 | 0.224 | 0.389 | 0.732 | 1.32 | 2.122 |
| CCNA_02934; conserved hypothetical protein | -0.003 | 0.633 | 1.144 | 2.146 | 2.127 | 2.572 |
| CCNA_03099; hypothetical protein | 0.002 | 0.378 | 0.678 | 1.271 | 1.525 | 2.473 |
| CCNA_03161; transcriptional regulator xylR | 0.017 | 1.201 | 1.769 | 2.792 | 2.9 | 2.983 |
| CCNA_03386; multimodular transpeptidase-transglycosylase PbpC | 0.002 | 0.775 | 1.546 | 2.13 | 2.477 | 2.602 |
| CCNA_03465; glycine cleavage system aminomethyltransferase T | 0.012 | 1.186 | 1.447 | 1.937 | 2.076 | 2.344 |
| CCNA_03611; glutathione-regulated potassium-efflux system protein kefC | 0.005 | -0.076 | 1.577 | 2.82 | 3.551 | 3.907 |
| CCNA_03803; acetyltransferase family holdfast biogenesis protein HfsK | -0.003 | 0.895 | 1.957 | 2.71 | 3.132 | 3.698 |
| CCNA_03902; conserved hypothetical protein | -0.009 | 0.681 | 1.566 | 2.037 | 2.339 | 3.165 |

**Table S6** *Strains and plasmids used in this study*

To analyze the sequence data used for fitness calculations, we used the *C. crescentus* NA1000 genome annotation. NA1000 is directly derived from CB15, but its genome has more detailed annotations and is better curated. To facilitate interpretation of the BarSeq data, genotypes in the text and supplemental tables use the NA1000 locus numbers and nomenclature. We note the corresponding CB15 locus numbers in the “Description” column.

| ***Strains*** |  |  |  | | |  |
| --- | --- | --- | --- | --- | --- | --- |
| **Strain name** | **Organism** | **Genotype** | **Description** | | | **Source** |
| FC19 | *C. crescentus* CB15 | CB15 | Wild-type | | | ATCC 19089 |
| FC1365 | *C. crescentus* CB15 | ∆*hfiA* | In-frame deletion of *CC_0817* | | | Ref. 10 |
| FC1974 | *C. crescentus* CB15 | ∆*hfsJ* | In-frame deletion of *CC_0095* | | | Ref. 10 |
| FC3105 | *C. crescentus* CB15 | ∆*pleD* | In-frame deletion of *CC_2462* | | | This work |
| FC3054 | *C. crescentus* CB15 | ∆*CCNA_00497* | In-frame deletion of *CC_0465* | | | This work |
| FC3055 | *C. crescentus* CB15 | ∆*CCNA_02386* | In-frame deletion of *CC_2301* | | | This work |
| FC3056 | *C. crescentus* CB15 | ∆*rfbB* | In-frame deletion of *CC_3629* | | | This work |
| FC3057 | *C. crescentus* CB15 | ∆*wbqP* | In-frame deletion of *CC_1486* | | | This work |
| FC3058 | *C. crescentus* CB15 | ∆*hfsJ* ∆*wbqP* | In-frame deletion of *CC_1486* in FC1974 background | | | This work |
| FC1266 | *C. crescentus* CB15 | ∆*flgH* | In-frame deletion of *CC_2066* | | | This work |
| FC3013 | *C. crescentus* CB15 | ∆*cpaH* | In-frame deletion of *CC_2940* | | | This work |
| FC1265 | *C. crescentus* CB15 | ∆*pilA* | In-frame deletion of *CC_2948* | | | This work |
| FC3019 | *C. crescentus* CB15 | ∆*CCNA_01242* | In-frame deletion of *CC_1184* | | | This work |
| FC3021 | *C. crescentus* CB15 | ∆*hfsL* | In-frame deletion of *CC_2277* | | | This work |
| FC3020 | *C. crescentus* CB15 | ∆*hfaE* | In-frame deletion of *CC_2639* | | | This work |
| FC3015 | *C. crescentus* CB15 | ∆*hfsJ* ∆*flgH* | In-frame deletion of *CC_2066* in FC1974 background | | | This work |
| FC3016 | *C. crescentus* CB15 | ∆*hfsJ* ∆*cpaH* | In-frame deletion of *CC_2940* in FC1974 background | | | This work |
| FC3107 | *C. crescentus* CB15 | ∆*flgH* ∆*cpaH* | In-frame deletion of *CC_2940* in FC1266 | | | This work |
| FC3085 | *C. crescentus* CB15 | ∆*flgH* ∆*hfiA* | In-frame deletion of *CC_0817* in FC1266 background | | | This work |
| FC3083 | *C. crescentus* CB15 | ∆*cpaH* ∆*hfiA* | In-frame deletion of *CC_0817* in FC3013 background | | | This work |
| FC3084 | *C. crescentus* CB15 | ∆*pilA* ∆*hfiA* | In-frame deletion of *CC_0817* in FC1265 background | | | This work |
| FC3104 | *C. crescentus* CB15 | ∆*flgH* ∆*pleD* | In-frame deletion of *CC_2462* in FC1266 background | | | This work |
| FC3103 | *C. crescentus* CB15 | ∆*cpaH* ∆*pleD* | In-frame deletion of *CC_2462* in FC3013 background | | | This work |
| FC3108 | *C. crescentus* CB15 | ∆*pilA* ∆*pleD* | In-frame deletion of *CC_2462* in FC1265 background | | | This work |
| FC3017 | *C. crescentus* CB15 | ∆*flgH* ∆*pilA* | In-frame deletion of *CC_2948* in FC1266 background | | | This work |
| FC3018 | *C. crescentus* CB15 | ∆*cpaH* ∆*pilA* | In-frame deletion of *CC_2948* in FC3013 background | | | This work |
| FC3097 | *C. crescentus* CB15 | ∆*CCNA_00497 xyl::P_xyl_-empty* | pXGFPC-2 integrated at xylose locus of FC3054 | | | This work |
| FC3098 | *C. crescentus* CB15 | ∆*CCNA_00497 xyl::P_xyl_-CCNA_00497* | pFC3080 integrated at xylose locus of FC3054 | | | This work |
| FC3101 | *C. crescentus* CB15 | ∆*CCNA_02386 xyl::P_xyl_-empty* | pXGFPC-2 integrated at xylose locus of FC3055 | | | This work |
| FC3102 | *C. crescentus* CB15 | ∆*CCNA_02386 xyl::P_xyl_-CCNA_02386* | pFC3082 integrated at xylose locus of FC3055 | | | This work |
| FC3095 | *C. crescentus* CB15 | ∆*rfbB xyl::P_xyl_-empty* | pXGFPC-2 integrated at xylose locus of FC3056 | | | This work |
| FC3096 | *C. crescentus* CB15 | ∆*rfbB xyl::P_xyl_-rfbB* | pFC3079 integrated at xylose locus of FC3056 | | | This work |
| FC3099 | *C. crescentus* CB15 | ∆*wbqP xyl::P_xyl_-empty* | pXGFPC-2 integrated at xylose locus of FC3057 | | | This work |
| FC3100 | *C. crescentus* CB15 | ∆*wbqP* *xyl::P_xyl_-wbqP* | pFC3081 integrated at xylose locus of FC3057 | | | This work |
| FC3075 | *C. crescentus* CB15 | ∆*flgH* *xyl::P_flgE_-empty* | pFC3094 integrated at xylose locus of FC1266 | | | This work |
| FC3074 | *C. crescentus* CB15 | ∆*flgH xyl::P_flgE_-flgH* | pFC3093 integrated at xylose locus of FC1266 | | | This work |
| FC3070 | *C. crescentus* CB15 | ∆*cpaH xyl::P_xyl_-empty* | pXGFPC-2 integrated at xylose locus of FC3013 | | | This work |
| FC3071 | *C. crescentus* CB15 | ∆*cpaH xyl::P_xyl_-cpaH* | pFC3090 integrated at xylose locus of FC3013 | | | This work |
| FC3073 | *C. crescentus* CB15 | ∆*pilA xyl::P_pilA_-empty* | pFC3092 integrated at xylose locus of FC1265 | | | This work |
| FC3072 | *C. crescentus* CB15 | ∆*pilA* *xyl::P_pilA_-pilA* | pFC3091 integrated at xylose locus of FC1265 | | | This work |
| FC3126 | *C. crescentus* CB15 | ∆*CCNA_01242 xyl::P_CCNA_01242_-empty* | pFC3124 integrated at xylose locus of FC3019 | | | This work |
| FC3127 | *C. crescentus* CB15 | ∆*CCNA_01242 xyl::P_CCNA_01242_-CCNA_01242* | pFC3125 integrated at xylose locus of FC3019 | | | This work |
| FC3088 | *C. crescentus* CB15 | ∆*hfsL xyl::P_xyl_-empty* | pXGFPC-2 integrated at xylose locus of FC3021 | | | This work |
| FC3089 | *C. crescentus* CB15 | ∆*hfsL* *xyl::P_xyl_-hfsL* | pFC3078 integrated at xylose locus of FC3021 | | | This work |
| FC3086 | *C. crescentus* CB15 | ∆*hfaE xyl::P_xyl_-empty* | pXGFPC-2 integrated at xylose locus of FC3020 | | | This work |
| FC3087 | *C. crescentus* CB15 | ∆*hfaE xyl::P_xyl_-hfaE* | pFC3077 integrated at xylose locus of FC3020 | | | This work |
| APA_752 | *E. coli* WM3064 | Tn-HiMar (Km^R^) | Bacroded transposon pool for created BarSeq libraries | | | Ref. 29 |
| ***Plasmids*** |  |  |  | | |  |
| **Plasmid name** | **Description** | | | **Antibiotic** | **Reference** | |
| pNPTS138 | Suicide plasmid for making unmarked deletions in *C. crescentus*; carries sacB for counter-selection | | | Km | M. R Alley  unpublished | |
| pFC3059 | To delete *CCNA_01242*; contains fusion of *CC_1184* flanking regions with first and last 12 nucleotides of *CC_1184* ORF included | | | Km | This work | |
| pFC3060 | To delete *hfaE*; contains fusion of *CC_2639* flanking regions with first and last 12 nucleotides of *CC_2639* ORF included | | | Km | This work | |
| pFC3061 | To delete *hfsL*; contains fusion of *CC_2277* flanking regions with first 12 and last 96 nucleotides of *CC_2277* ORF included | | | Km | This work | |
| pFC3063 | To delete *CCNA_00497*; contains fusion of *CC_0465* flanking regions with first and last 12 nucleotides of *CC_0465* ORF included | | | Km | This work | |
| pFC3062 | To delete *CCNA_02386*; contains fusion of *CC_2301* flanking regions with first and last 12 nucleotides of *CC_2301* ORF included | | | Km | This work | |
| pFC3068 | To delete *rfbB*; contains fusion of *CC_3629* flanking regions with first and last 12 nucleotides of *CC_3629* ORF included | | | Km | This work | |
| pFC3069 | To delete *wbqP*; contains fusion of *CC_1486* flanking regions with first and last 12 nucleotides of *CC_1486* ORF included | | | Km | This work | |
| pFC1267 | To delete *pilA*; contains fusion of *CC_2948* flanking regions with first and last 12 nucleotides of *CC_2948* ORF included | | | Km | This work | |
| pFC1268 | To delete *flgH*; contains fusion of *CC_2066* flanking regions with first and last 12 nucleotides of *CC_2066* ORF included | | | Km | This work | |
| pFC3065 | To delete *cpaH*; contains fusion of *CC_2940* flanking regions with first and last 12 nucleotides of *CC_2940* ORF included | | | Km | This work | |
| pFC3067 | To delete *pleD*; contains fusion of *CC_2462* flanking regions with first and last 12 nucleotides of*CC_2462* ORF included | | | Km | This work | |
| pXGFPC-2  (pMT585) | Contains multiple cloning site downstream of Pxyl; integrates upstream of *xylX*; used for complementations | | | Km | Ref. 50 | |
| pFC3077 | pXGFPC-2 containing *hfaE* under the control of *P_xyl_* for integration at *xylX* locus | | | Km | This work | |
| pFC3078 | pXGFPC-2 containing *hfsL* under the control of *P_xyl_* for integration at *xylX* locus | | | Km | This work | |
| pFC3080 | pXGFPC-2 containing *CCNA_00497* under the control of *P_xyl_* for integration at *xylX* locus | | | Km | This work | |
| pFC3082 | pXGFPC-2 containing *CCNA_02386* under the control of *P_xyl_* for integration at *xylX* locus | | | Km | This work | |
| pFC3079 | pXGFPC-2 containing *rfbB* under the control of *P_xyl_* for integration at *xylX* locus | | | Km | This work | |
| pFC3081 | pXGFPC-2 containing *wbqP* under the control of *P_xyl_* for integration at *xylX* locus | | | Km | This work | |
| pFC3091 | pXGFPC-2 containing *pilA* under the control of *P_pilA_* for integration at *xylX* locus; 316bp upstream of *CC_2948* fused to the *CC_2948* ORF was inserted in reverse oreintation into pXGFPC-2 | | | Km | This work | |
| pFC3092 | pXGFPC-2 containing *P_pilA_* without insert for integration at *xylX* locus; 316bp upstream of *CC_2948* was inserted in reverse orientation into pXGFPC-2 | | | Km | This work | |
| pFC3093 | pXGFPC-2 containing *flgH* under the control of *P_flgF_* for integration at *xylX* locus; 226bp upstream of *CC_2063* fused to the *CC_2066* ORF was inserted in reverse oreintation into pXGFPC-2 | | | Km | This work | |
| pFC3094 | pXGFPC-2 containing *P_flgE_* without insert for integration at *xylX* locus; 216bp upstream of *CC_2063* was inserted in reverse orientation into pXGFPC-2 | | | Km | This work | |
| pFC3090 | pXGFPC-2 containing *cpaH* under the control of *P_xyl_* for integration at *xylX* locus | | | Km | This work | |
| pFC3124 | pXGFPC-2 containing *P_CCNA_01242_* without insert for integration at *xylX* locus; 99bp upstream of the *CC_1184* ORF was inserted in reverse orientation into pXGFPC-2 | | | Km | This work | |
| pFC3124 | pXGFPC-2 containing *CCNA_01242* under the control of *P_CCNA_01242_* for integration at *xylX* locus; 99bp upstream of *CC_1184* fused to the *CC_1184* ORF was inserted in reverse oreintation into pXGFPC-2 | | | Km | This work | |
| pFC1948 | pRKlac290 containing the *hfiA* promoter fused to *lacZ* | | | Tet | Ref. 10 | |

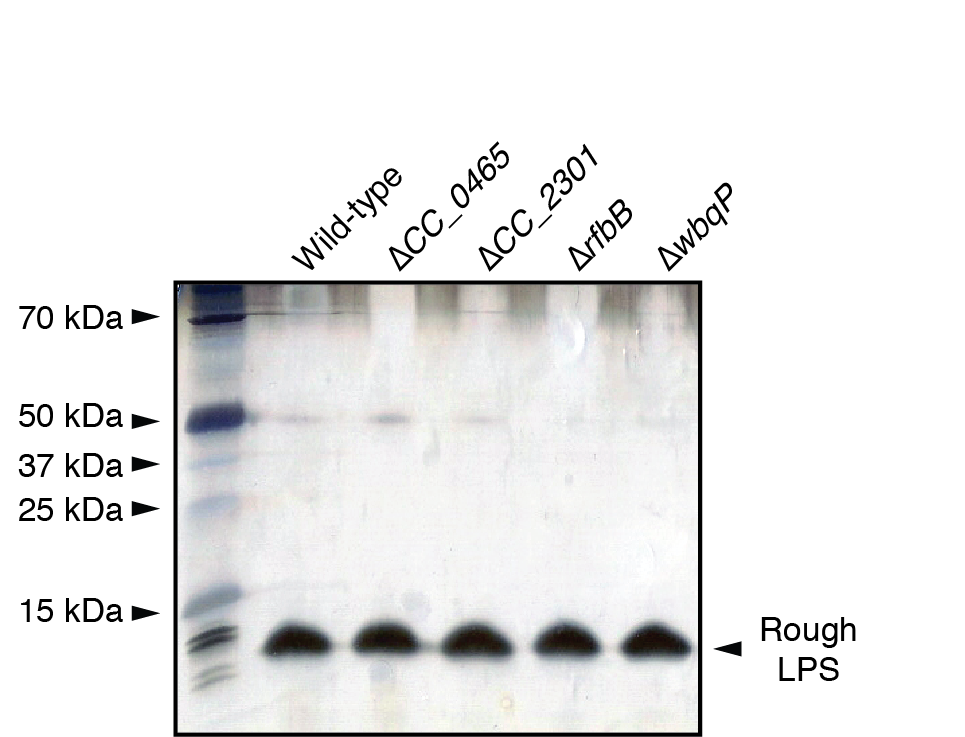

**Figure S1** *Analysis of rough LPS in SLPS mutants*

Silver stained polyacrylamide gel of rough LPS was isolated and analyzed as described in materials and methods. None of the four SLPS mutants show apparent defects in rough LPS.

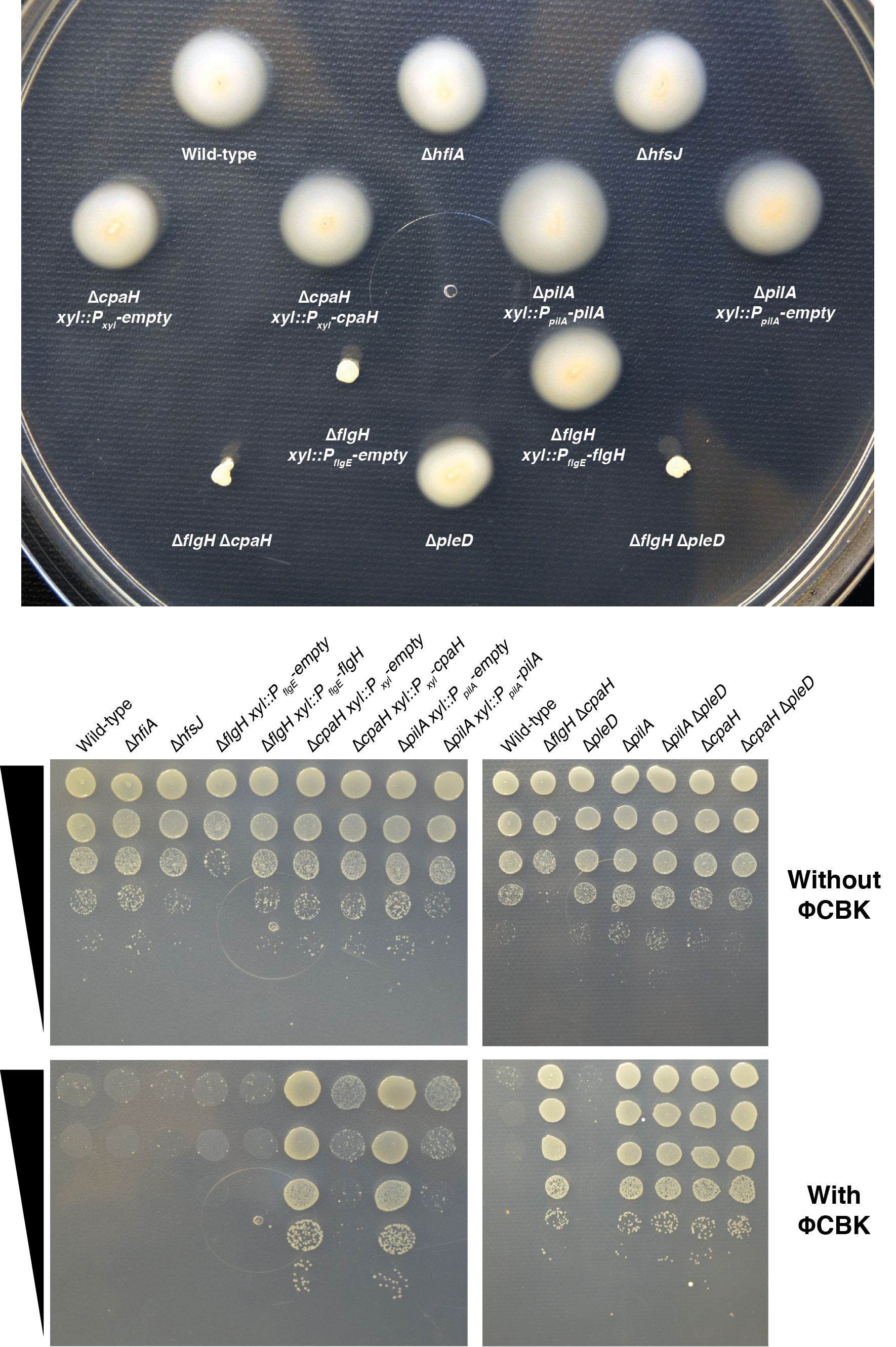
**­­­**

**Figure S2** *Validation of pilus and flagellum phenotypes in polar appendage mutants*

Top: Swimming phenotypes for relevant mutants in swarm agar. Note that ∆*pilA* has a slight, but reproducible, increase in swarm size that can be complemented by ectopic expression of *pilA* in trans. Mutants lacking *flgH* show the expected non-motile phenotype. The data are representative of four independent experiments. Bottom: ΦCBK sensitivity of polar appendage mutants. Mutants lacking *pilA* or *cpaH* show the expected ΦCBK resistance phenotype. The data are representative of three independent experiments.

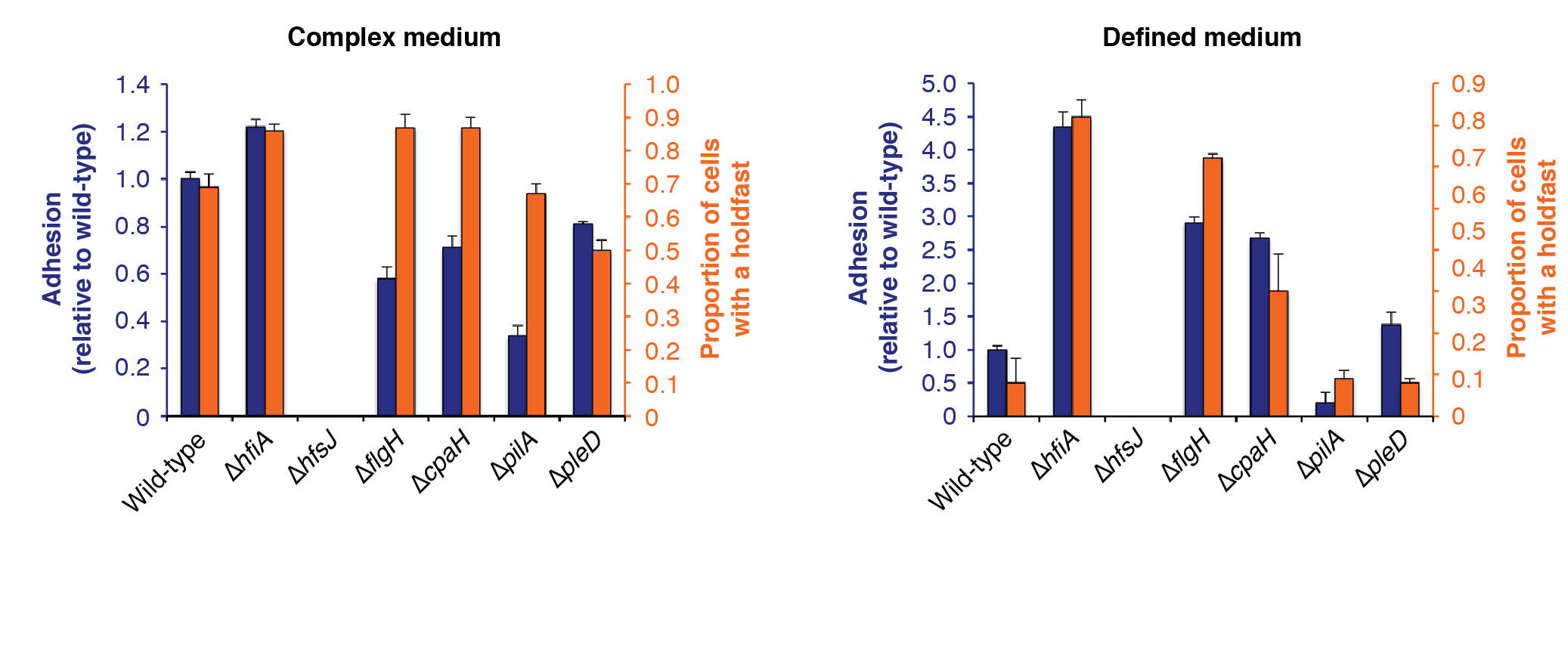

**Figure S3** *Comparison of crystal violet staining and holdfast counts for polar appendage mutants*

Surface attachment was assessed by CV staining (violet bars) and holdfast production by fWGA staining (orange bars) as described in Material and Methods. The values for growth in complex medium are shown on the left and defined medium on the right.

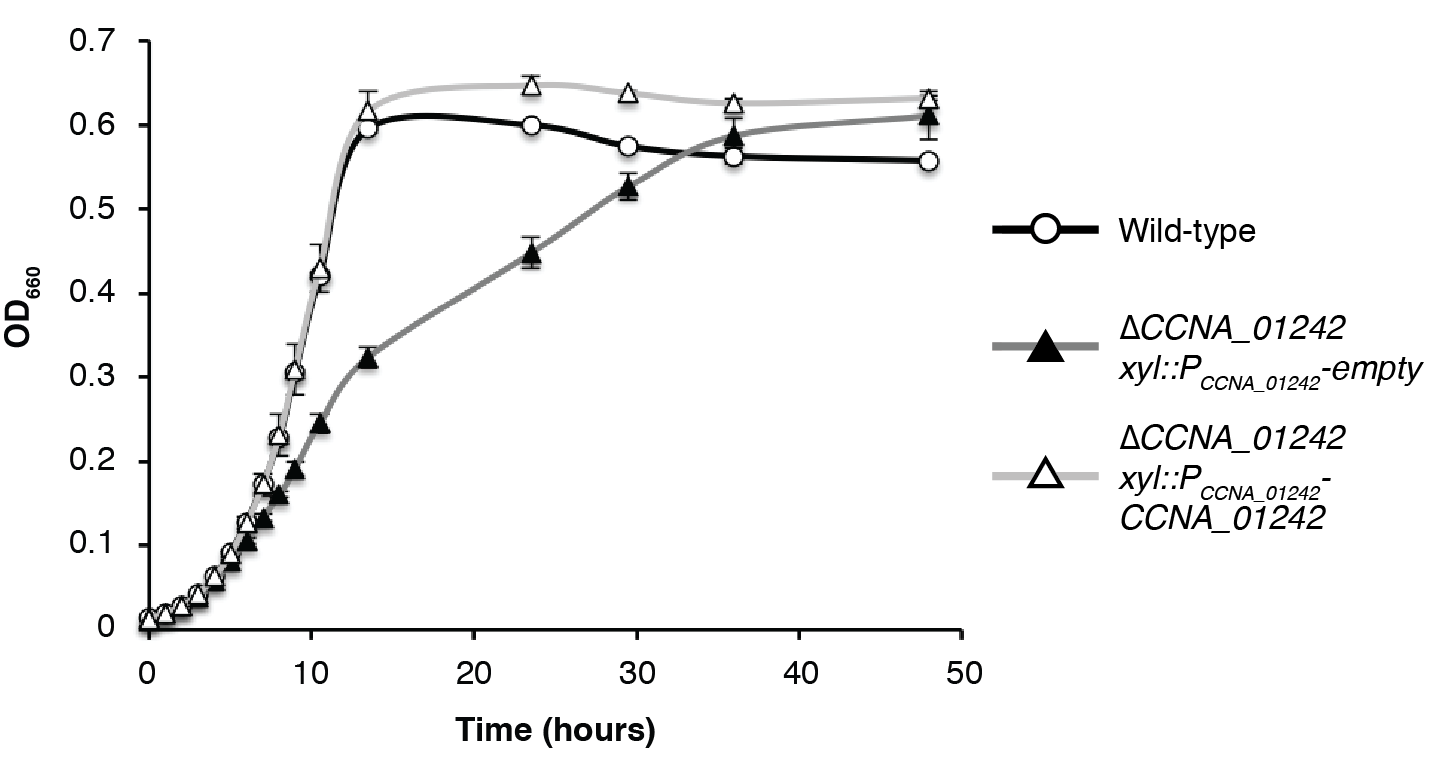

**Figure S4** *Complementation of growth defect in ∆*CCNA_01242

The ∆*CCNA_01242* mutant displays a biphasic growth curve indicating a defect in later growth phases. Normal growth is restored by ectopic complementation with *CCNA_01242.*
